# Network Thermodynamics of Biological Systems: A Bond Graph Approach

**DOI:** 10.1101/2022.05.04.490650

**Authors:** Peter J. Gawthrop, Michael Pan

## Abstract

Edmund Crampin (1973-2021) was at the forefront of Systems Biology research and his work will influence the field for years to come. This paper brings together and summarises the seminal work of his group in applying energy-based bond graph methods to biological systems. In particular, this paper: (a) motivates the need to consider energy in modelling biology; (b) introduces bond graphs as a methodology for achieving this; (c) describes extensions to modelling electrochemical transduction; (d) outlines how bond graph models can be constructed in a modular manner and (e) describes stoichiometric approaches to deriving fundamental properties of reaction networks. These concepts are illustrated using a new bond graph model of photosynthesis in chloroplasts.

## 1. Introduction

Edmund Crampin [1] made seminal contributions to Systems Biology which will remain influential in the field. One of his many research areas was the energy-based analysis of biological systems using the bond graph approach and he coauthored an number of papers [2–17] and directly influenced others [18–23]. This paper brings together and summarises this work, illustrates the approach using photosynthesis as an example and suggests future research directions.

One of the great challenges in biology is understanding how the complex nature of biochemistry leads to robust physiological function [24]. Mathematical models (and mechanistic models in particular) are essential to both bridging between the vast range of biological measurements and making sense of how biological systems function. However, biochemical systems are highly diverse both in their physics and in how they operate, making it difficult for a single modelling approach to consistently capture all aspects of biology [25]. Nonetheless, underlying all biological systems are the laws of physics, particularly conservation of mass, charge, momentum and energy. Energy in particular is central to life, just as it is central to physics: it can be take different forms but can never be created or destroyed. Thus, models that explicitly consider energy can connect and communicate with each other through this conserved physical quantity (Figure 1). Bond graphs build on this insight, combining energy transduction with the dynamics of biological systems.

**Figure 1:**
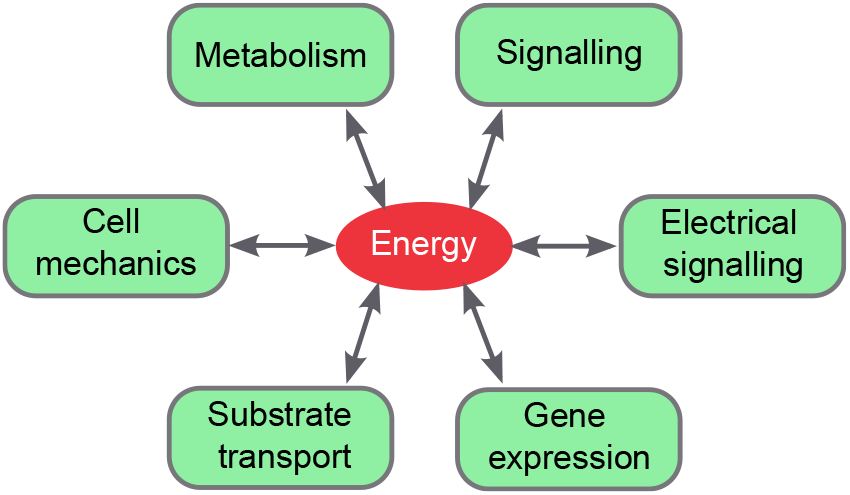
Energy as a common currency of biological systems. Cellular biochemistry comprises a diverse range of subsystems spanning across different physical domains. However, as with all physical systems, they share energy in common. Explicitly modelling energy allows for disparate cellular processes to be linked together.

Bond graphs were introduced by Paynter [26, 27] to model the flow of *energy* though physical systems of interest to engineers and are described in several text books [28–31] and a tutorial for control engineers [32]. Bond graphs provide a systematic approach to elucidating the *analogies* between disparate physical domains based on energy. They are related to the behavioural approach to systems dynamics [33] and to the port-Hamiltonian [34] methodology. The basic principles of the bond graph approach are discussed in § 2.

Bond graphs were first used to model chemical reaction networks by Katchalsky and coworkers [35]. A detailed account is given by Oster et al. [36] and an overview of the approach is given by Perelson [37]. Katchalsky’s work was further developed in the context of chemical reaction systems by a number of groups [38–43]. Later, this work was extended in the context of Systems Biology by Edmund Crampin’s group at Melbourne. These extensions are discussed in § 3 – § 7.

Bond graphs represent the flow of energy though systems and, as such, can be used to represent and connect systems in more than one physical domain; thus, for example, the transduction of chemical energy and electrical energy can be readily modelled within this framework [5, 10, 19, 22]. Chemoelectrical transduction is discussed in § 4 in the context of redox reactions.

Modelling complex biological systems requires complex modelling problems to be broken down into manageable pieces. This can be enabled by developing models of subsystems that are subsequently coupled together in a hierarchical manner. Because they are based on a graph structure, bond graphs are relatively simple to merge, and the underlying physical representation ensures that merged models remain consistent with the laws of physics [13–16, 21]. In § 5, we discuss how bond graphs can be embedded within reusable modules. This process is facilitated by a set of Python-based tools `BondGraphTools` [13]. § 6 discusses the relationship beween bond graphs, stoichiometric analysis, pathway analysis and CRNs.

Photosynthesis [44–47] is an energy transduction system converting the energy of photons to chemical energy stored as ATP via the transduction of the energy of light, chemicals, electrons and protons. A simple bond graph model of photosynthesis is given in § 7 and used throughout the paper to provide illustrative examples.

§ 8 gives directions for future research.

## 2. Energy-based analogies

In the 19th century, James Clerk Maxwell observed [48] that analogies are central to scientific thinking and allow mathematical results and intuition from one physical domain (e.g. electrical, mechanical or chemical) to be transferred to another. Bond-graph modeling is based on three types of analogies, namely, *variable* analogies, *component* analogies, and *connection* analogies [32]. These analogies arise in the essential elements of a bond graph: bonds (§ 2.1), components (§ 2.2) and junctions (§ 2.3).

### 2.1. Variable analogies and bonds

The bond graph approach of Paynter [26, 27] uses the effort-flow analogy of Table 1. In particular, Table 1 shows three categories of variable (effort, flow and quantity) with examples from two physical domains (electrical and chemical):

**Effort** variables, with the generic symbol *e*, including mechanical force, electrical voltage and Gibbs energy.
**Flow** variables, with the generic symbol *f*, including mechanical velocity, electrical current and molar flow.
**Quantity** variables, with the generic symbol *q,* including electrical charge and mechanical displacement. These are the integral with re**s**pect to time of the correponding flows: *q* = *∫*^*t*^ (*τ*)*dτ*.

**Table 1:** Analogous variables. Systematic modeling, including the bond graph approach, uses the concept of analogous variables to bring together different physical domains. One such analogy is the effort/flow analogy displayed here: each row contains analogous variables, each column corresponds to a domain. In each case effort × flow = power. Gibbs energy is also referred to as *chemical potential.*

| Bond-Graph | Electrical | Chemical |
| --- | --- | --- |
| Effort<br>$e$ | Voltage<br>$V$ (V) | Gibbs energy<br>$\mu$ (J mol<sup>-1</sup>) |
| Flow<br>$f$ | Current<br>$I$ (A) | Molar flow<br>$v$ (mol/sec) |
| Quantity<br>$q = \int^t f(\tau) d\tau$ | Charge<br>$q$ (C) | Molar amount<br>$x$ (mol) |

Additional domains, including mechanical, magnetic and thermal, can also be incorporated in this scheme. We note that the quantity variables are linked to effort and flows: effort has unit J/quantity and flow has unit quantity/second. Thus, a key insight is that the product of the effort and flow variables in each domain is power, that is,

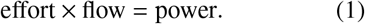

Hence, different physical domains, despite disparateeffort and flow variables, have *power* in common. This insight leads to the definition of the power *bond* which carries effort and flow variables; in bond graph notation it has the symbol 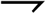.

In the context of biochemical systems, models have a long history of incorporating flow. The effort (chemical potential) has historically been neglected, but has been considered more regularly in light of new thermodynamic measurements [49] and modelling studies using thermodynamics to restrict the range of plausible parameters [50–52].

### 2.2. Component analogies and components

In addition to variable analogies, physical domains often share similar mathematical equations. These are encapsulated via component analogies. Some components common to several physical domains include the following:

**R** component, which can correspond to an electrical resistor or a mechanical damper, *dissipates* energy;
**C** component, which can correspond to an electrical capacitor or a mechanical spring (or compliance), *stores* energy through its charge *q* = ∫ *f dt*;
**I** component, which can correspond to an electrical inductor or a mechanical mass, *stores* energy through its generalised momentum *p = ∫ edt.*

In the linear case the corresponding equations for the **R**, **C**, and **I** components in terms of the generic variables of Table 1 are, respectively,

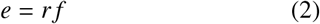

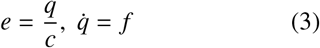

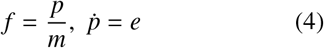

where *r, c,* and *m* are constants describing the corresponding physical system. In the electrical case, (2) corresponds to Ohm’s law and (3) to Coulomb’s law; in the mechanical case, (3) corresponds to Hooke’s law, while (4) corresponds to Newton’s second law.

### 2.3. Connection analogies and junctions

Electrical components may be connected in *parallel* (where the *voltage* is common) and *series* (where the *current* is common). These two concepts are generalised in the bond graph notation as the **0** and **1** junctions respectively. The **0** junction of Figure 2(a) implies that all impinging bonds have the same *potential* (but different flows). The **1** junction of Figure 2(b) implies that all impinging bonds have the same *flow* (but different potentials). The direction of positive energy transmission is determined by the bond half arrow. As all bonds impinging on a **0** junction have the same *potential,* the half arrow implies the sign of the *flows* for each impinging bond. The reverse is true for **1** junctions, where the half arrow implies the signs of the *potentials*.

**Figure 2:**
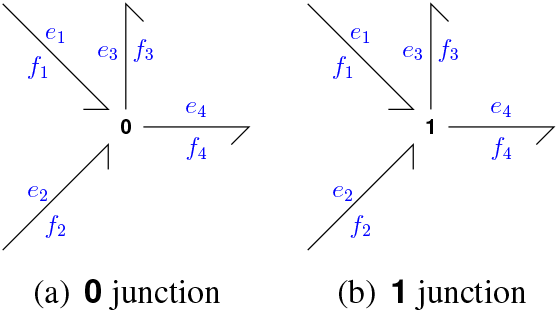
**1** and **0** junctions

To be explicit, Figure 2(a) shows a **0** junction with four impinging bonds. The *potentials* on the four bonds are equal and the four flows are related by a single equation having regard to the half arrows:

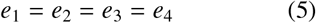

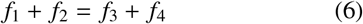

Similarly, Figure 2(b) implies that

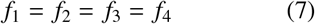

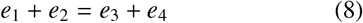

These junctions therefore describe the *network topology* of the system.

## 3. Chemical Reactions

Like all physical systems, biological systems must comply with conservation of energy. Energy is at the core of how biological systems operate and has been proposed to have lead to the origins of complex life [53]. It follows that understanding living systems requires a modelling approach that satisfies energy conservation. Bond graphs satisfy this requirement, as we discuss in this section. § 3.1 considers a simple reaction, §§ 3.2 and 3.3 introduces the thermodynamics of species and reaction components and § 3.4 shows how reactions can be connected to form a network. § 3.5 looks at multiple stoichiometry and § 3.6 details a diagrammatic simplification of reaction components.

### 3.1. A simple reaction 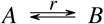

The simple reaction 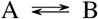 is shown as a bond graph in Figure 3. This bond graph contains two *bonds* 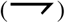 and three *components* **Ce**:**A**, **Re**:**r** and **Ce**:**B**. Note that bond graph components follow the convention of ***type***:*name*, i.e. the component type (e.g. **Ce**) is separated from the component name (e.g. **A**) by a colon.

**Figure 3:**
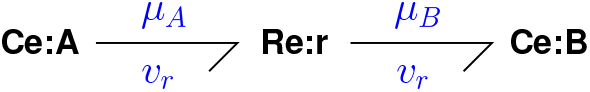
Bond graph of the reaction 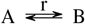. *μ_A_* is the chemical potential associated with the chemical species **Ce**:**A**, *μ*_B_ is the chemical potential associated with **Ce**:**B** and *v_r_* is the molar flow associated with the reaction **Re**:**r**.

The bond graph is to be interpreted as follows:

**Ce** components represent chemical species. In particular **Ce**:**A** and **Ce**:**B** represent the two chemical species A and B respectively. They are special cases of the bond graph **C** component which *store chemical energy* in the same way as electrical capacitors store electrical energy.
**Re** components represent *reversible* chemical reactions between species. The inward impinging bond is associated with reactants and the outward impinging bond is associated with reaction products. In particular, **Re**:**r** represents the chemical reaction between the two species A and B where A is the reactant and B the product. It is a special case of the bond graph **R** component which *dissipates* chemical energy. It is important to realise that calling A and B reactant and product respectively is a merely conventional designation. The reaction can proceed in either direction and is exactly equivalent to the reaction 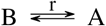 where the reactant and product designation is reversed. The **Re** component is discussed in detail in §3.3,p.5.
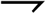 The components are connected by *bonds* (in the bond graph sense of the word) represented by 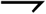. The bonds *transmit* chemical energy and are associated with a *reaction flow* v and a *chemical potential* μ. (The annotation with potentials *μ* and flows *v* is for clarity and does not form part of the bond graph). The half-arrow on the bond pointing into the **Re** component indicates that species A is to appear on the left-hand side of the reaction; the half-arrow on the bond pointing out of the **Re** component indicates that species B is to appear on the right-hand side of the reaction. The half-arrow does *not* represent direction of flow; this is a reversible reaction and the flow can be in either direction. It does, however, indicate that a reaction flow from left to right is regarded as positive and that from right to left as negative.

### 3.2. The Bond Graph **Ce** component

The **C** component is the bond graph abstraction of an electrical capacitor and is used as such in the context of electrochemistry in § 4. For the purposes of modelling chemical species, it is convenient to define a modified nonlinear **C** component: the **Ce** component.

Thus, the **Ce** component represents a chemical species with chemical potential replacing voltage and molar flow replacing current [35, 36]. In solutions with constant temperature and pressure, the Gibbs free energy determines the chemical potential of species [54]. Thus, the bond graph **Ce** component for biomolecular systems accumulates a chemical species A as the number of moles *x_A_* and generates the corresponding chemical potential *μ_A_* in terms of the molar flow *x_A_* [2]. In dilute solutions, the following relationships hold:

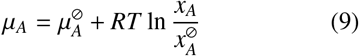

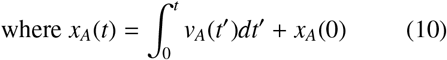

where 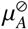 is the chemical potential of *x_A_* when 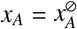. When thermodynamic measurements are available, 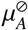 is often set to the standard free energy of formation and the standard amount is set to *x_A_* = *c*_0_W, where *c*_0_ is the standard concentration (often 1 M) and W [litres] is the compartment volume. Equation (9) may be rewritten in two ways:

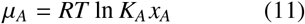

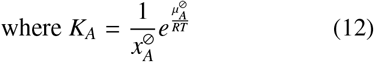

and

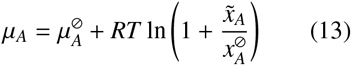

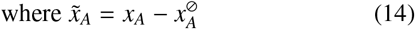

Equation (12) is equivalent to that used previously [2] and equation (14) is convenient when 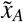 is small and so:

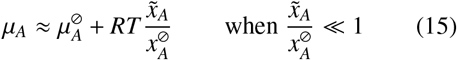

### 3.3. The Bond Graph **Re** component

The **R** component is the bond graph abstraction of an electrical resistor. In the chemical context, a two-port **R** component represents a chemical reaction with net chemical potential replacing voltage and molar flow replacing current [35, 36]. As it is so fundamental, this two port **R** component is given a special symbol: **Re** [2]. The **Re** component determines a reaction flow v in terms of forward and reverse affinities *A^f^* and *A^r^* as the *Marcelin–de Donder* formula [55]:

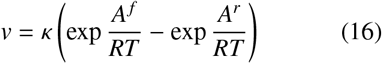

where *κ* is a rate constant, *A^f^* is the sum of chemical potentials in the reactants and *A^r^* is the sum of chemical potentials in the reactants. In the special case of mass-action kinetics, *κ* is a constant. In more general cases, *κ* may be a function of the forward and reverse affinities *A^f^* and *A^r^* or even the individual chemical potentials *μ* of each reactant and product [14, 15].

Thus, for the bond graph in Figure 3, the chemical potentials are given by

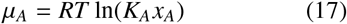

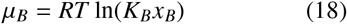

and the reaction rate is given by

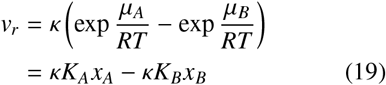

where the second equality arises from substituting Equations (17)–(18). Thus, the differential equations of the system are

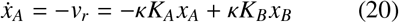

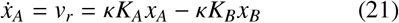

These are equivalent to mass action kinetics, but formulated using the thermodynamic parameters *κ* and *K.* This new parameterisation ensures that models are thermodynamically consistent, which kinetic parameters often fail to satisfy [10, 13].

### 3.4. Reactions with connections

Bond graphs can represent reactions more complex than the reaction 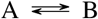 discussed in § 3.1. These require the bond graph ***0*** and ***1*** junction components of § 2.3.

When a reaction involves more than one reactant or product, **1** junctions are used. For example, Figure 4(a) is the bond graph representation of the reaction 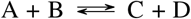. The left-hand **1** junction enforces two constraints:

1. The flow associated with each of the three bonds is the same (*v*).
2. To ensure chemical potential is conserved, the effort (forward chemical potential *A^f^*) associated with the bond connecting the **1** junction to the **Re** component is the *sum* of the chemical potentials *μ_A_* and *μ_B_* associated with species A and B:

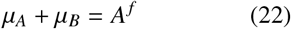 Half-arrows pointing *into* the junction correspond to the left-hand side of the equation and half-arrows pointing *out of* the junction correspond to the right-hand side of the equation. The right-hand **1** junction is similar, but now the effort (reverse chemical potential *A^r^*) associated with the bond connecting the **1** junction to the **Re** component is:

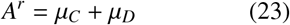

**Figure 4:**
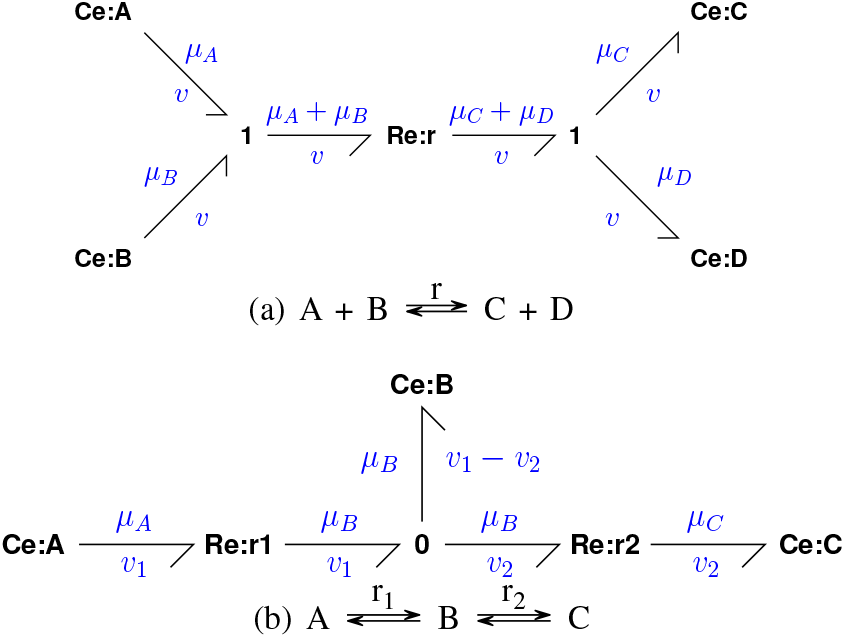
Bond graphs of reactions with connections.

When a species is involved in multiple reactions, **0** junctions are used. For example, the reaction 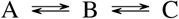 is represented by the bond graph in Figure 4(b), where the **0** junction enforces two constraints:

1. The effort (chemical potential *μ_B_*) associated with each of the three bonds is the same.
2. To ensure mass conservation, the flow into **Ce**:**B** is the *difference* between the flows associated with **Re**:**r1** and **Re**:**r2**:

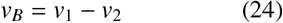 Again, half-arrows pointing *into* the junction correspond to the left-hand side of the equation; half-arrows pointing *out of* the junction correspond to the right-hand side of the equation.

### 3.5. Multiple stoichiometry

In some reactions, more than one molecule of a species appears. There are two ways of representing multiple molecules using bond graphs

1. Using multiple bonds [19].
2. Using the transformer (**TF**) component [2,35–41].

In this section, the multiple bond approach is used, but the **TF** approach is used in § 6.

Figure 5(a) shows a simple example of the reaction 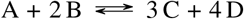. Here, the reaction affinities are given by

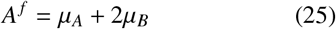

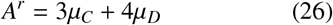

and the differential equations are given by 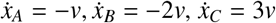 and 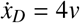, where *v* is the rate of reaction.

**Figure 5:**
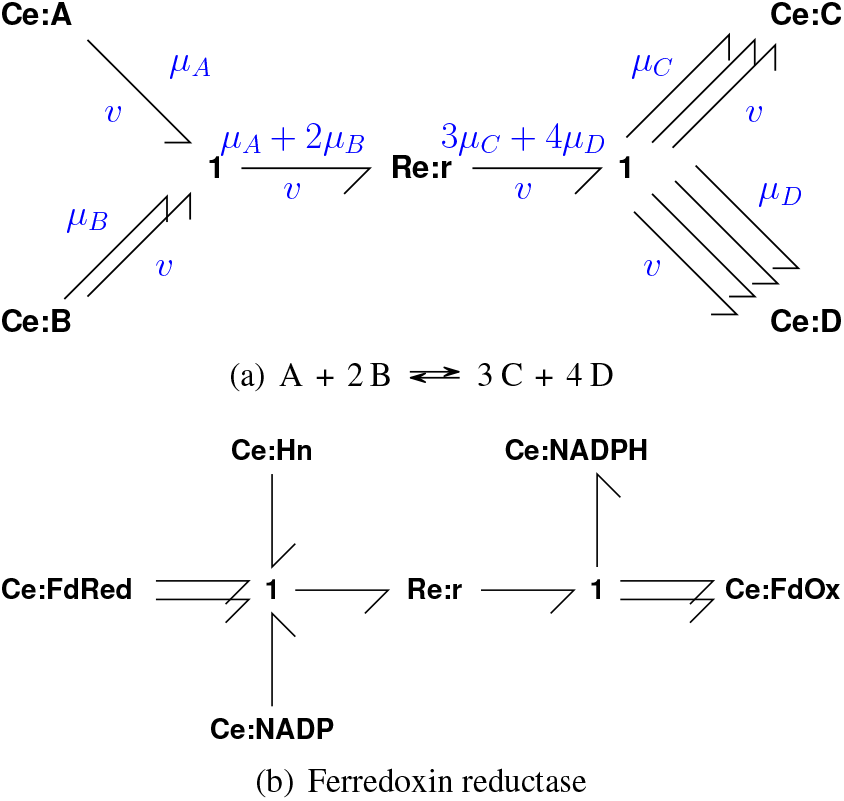
Bond graphs of reactions involving multiple stoichiometry.

Figure 5(b) shows the ferredoxin redox reaction which occurs in the photosynthesis electron transport chain [44] and is discussed in § 4.2 and § 7:

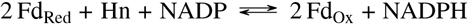

### 3.6. One-port **Re** component

While the two-port **Re** component is the most precise way of representing a reaction, one can also employ a shorthand to replace it with an **R** component connected to a **1** junction. For example, the bond graphs in Figures 6(a) and 6(b) are shorthands for those in Figures 5(a) and 5(b).

**Figure 6:**
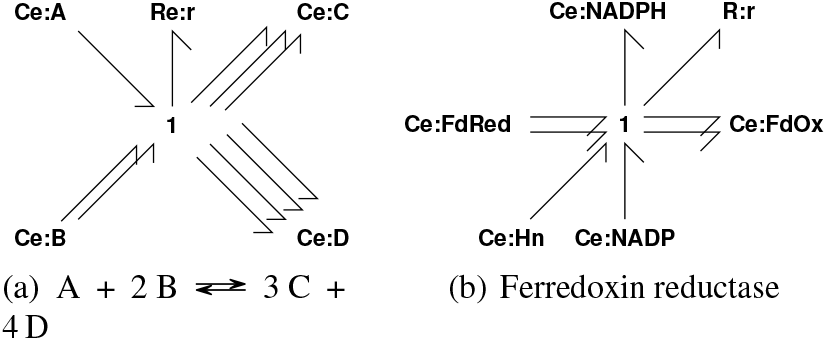
One-port **Re** component. Figures 6(a) and 6(b) are one-port **Re** representations of Figures 5(a) and 5(b) respectively.

This has two interpretations:

1. Since the shorthand representation preserves the reaction thermodynamics and stoichiometry, it can serve as a representation of the model structure in the absence of kinetics or regulation. Here, one can think of kinetics and regulation as modulators of the resistance parameter of the **R** component.
2. Under the assumption that all reactions follow the law of mass action with positive stoichiometries, there is a one-to-one mapping between the shorthand and full bond graph. Thus, the full bond graph can be uniquely generated from the shorthand representation.

In some cases, the one-port representation provides a clearer representation of the conservation laws of a biochemical network: this is used in § 4.2 and § 7. However, in general, the two-port **Re** component is used to give a mathematically explicit representation.

## 4. Electrochemical transduction

The fundamental biophysical processes of life involve the transduction of chemical energy and electrical energy. To provide two examples, the chemiosmotic theory of Mitchell [56] explains how both chemical and electrical energy are stored in a trans-membrane proton gradient; and Hodgkin and Huxley [57] show how the mutual transduction of chemical and electrical energy gives rise to action potential in nerves. Because the chemical and electrical domains are so intertwined, the analysis and understanding of such systems is enhanced by a common approach to the two domains. One example of this is the *proton motive force* PMF of chemiosmotic theory [45, 47, 58] which reexpresses the chemical potential of protons as electrical voltage using the Faraday constant so that it can be added to the electrical potential. A second example is the notion of *redox potential* [47, 59] which assigns a voltage to reactions involving electron transfer.

A bond graph interpretation of chemiosmosis [19] and membrane transduction [5, 10] has been developed. A general discussion of bioelectrical systems in a bond graph context is given by Gawthrop and Pan [22]. The notion of reexpressing chemical potential as electrical potential is not just confined to electrically-charged ions, but can be generally applied to any chemical species – charged or not [19]. This *Faraday-Equivalent Potential* is examined in § 4.1 and its implications for redox reactions are discussed in § 4.2.

### 4.1. Faraday-Equivalent Potential

The conversion factor relating the quantities charge and molar amount (see Table 1) from the electrical and chemical domains is *Faraday’s constant*

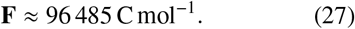

Noting that the units of electrical current are C s^-1^ or A and that the units of electrical voltage are J C^-1^ or V. Then the chemical potential μ [J mol^-1^] can be reexpressed as *ϕ* [V] and flow *v* [mol s^-1^] can be reexpressed as *f* [A] where:

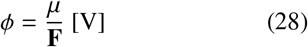

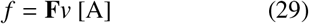

For example, consider NAD at standard conditions which has a chemical potential of 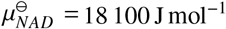 under standard conditions. The corresponding Faraday-equivalent potential is 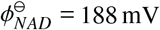. Similarly, a molar flow of *v* = 1 μmol s^-1^ has a Faraday-equivalent flow of about *f* = 97 mA.

### 4.2. Redox reactions

Redox reactions provide the energy required to sustain life [59, 60] and the notion of the *redox potential* is useful in describing their energetic properties. Both redox reactions and redox potential can be clearly and explicitly described using the bond graph approach. *Redox reactions* can be rewritten as two *half reactions* which explicitly account for electron transfer [47, 59]. Such reactions have a bond graph representation [19]. For example, the ferredoxin reductase reaction (Figure 5(b)) can be rewritten in terms of two half reactions r_1_ and r_3_ and an electron e^-^ transfer reaction r_2_:

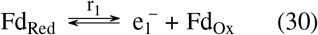

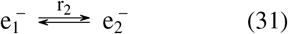

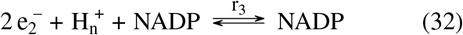

where 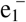 denotes electrons donated in half-reaction r_1_ (30) and 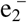 denotes electrons accepted in half-reaction r_3_ (32).

These half-reactions correspond to the bond graph representation of Figure 7. Thus, the bond graph component **Re**:**r1**, together with the components **C**:**FDred**, **C**:**FDox** and connecting bonds represents reaction r_1_ and the bond graph component **Re**:**r3**, together with the components **C**:**NADP**, **C**:**NADPH** and **C**:**Hn** and connecting bonds represents reaction r_3_.

**Figure 7:**
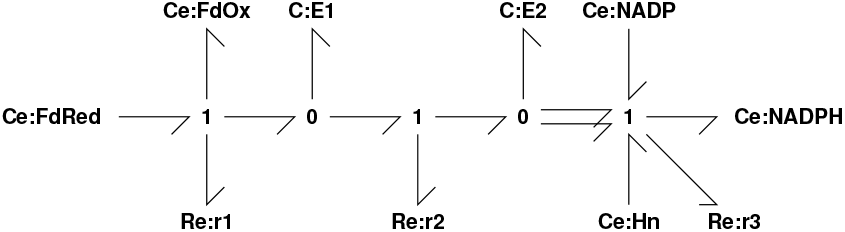
Ferredoxin reductase (Figure 5(b)) in redox form. Here, the overall redox reaction has been decomposed into two halfreactions (**Re**:**rl** and **Re**:**r3**) and an electron transfer reaction (**Re**:**r2**).

The ferredoxin reductase reaction in the redox form of Figure 7 is used as a module in a modular description of photosynthesis in § 7.

## 5. Modularity

By its very nature, systems biology is complex. Constructing large-scale models requires smaller ones to be composed together. A common challenge is finding interfaces for models to communicate. Bond graphs provide a natural interface for this: connections correspond to conservation laws in physics (§ 2) [61]. Here we discuss two methodologies for connecting models.

### 5.1. Black-box modularity

Traditionally, bond graphs define modules through a black-box paradigm that follows in the tradition of engineering. Here, modules are independently developed bond graph models with predefined interfaces to the external environment. This is useful in cases where connections are known in advance, for example, enzymes in a metabolic network [3].

Consider the enzyme-catalysed reaction

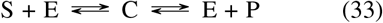

In this network, we consider the substrates S and P to be external since they interact with other enzymes. The enzyme states E and C are assumed to be internal since they are isolated in the context of metabolic networks (although this assumption would need to be relaxed when enzyme abundances are regulated through gene expression). The corresponding bond graph is in Figure 8(a), where the connection to the **Ce**:**S** and **Ce**:**P** components are left unconnected, like an open port in an electrical circuit. In bond graph notation, these are represented by **SS** components (**SS**:**S** and **SS**:**P**) that label the external connections.

**Figure 8:**
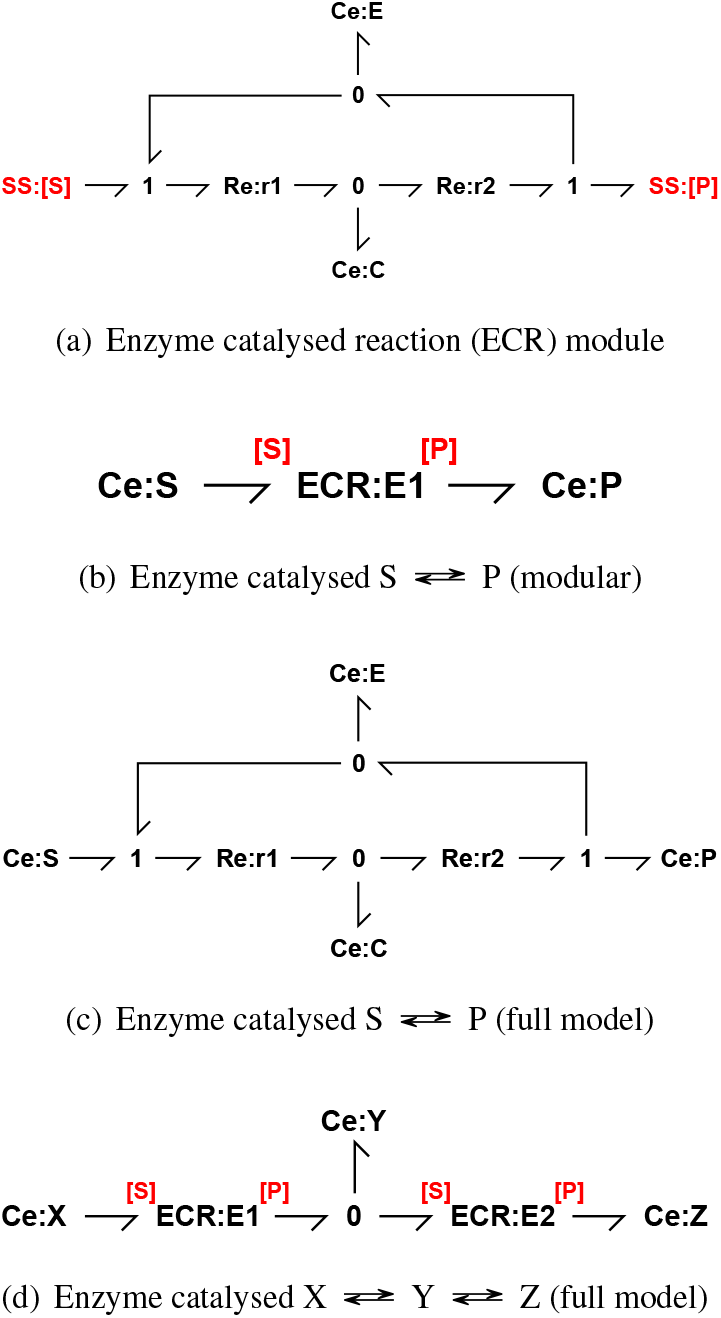
Modular representation of an enzyme catalysed reaction. (a) A modular representation of an enzyme-catalysed reaction (ECR). Here, the substrate S and product P are considered external, and are represented as external ports (**SS** components). This defines the reusable **ECR** module. (b) An enzyme catalysed reaction 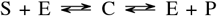 that uses the **ECR** module in (a). (c) The full representation of the bond graph in (b). (d) A bond graph representation of a series of enzyme catalysed reactions 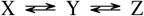, where each reaction is catalysed by the **ECR** module in (b).

We define the enzyme module in Figure 8(a) to be an **ECR** module. This can be reused within an outer bond graph to construct a model of the full reaction in Equation (33). This is shown in Figure 8(b). Because the **ECR** module has two ports, the ports need to be explicitly specified so that the components are connected correctly. These ports are labelled using the red [S] and [P] labels, indicating that the **Ce**:**S** component is connected to the [S] port and the **Ce**:**P** component is connected to the [P] port. An equivalent full bond graph representation is shown in Figure 8(c).

Another advantage of this modular representation is that the enzyme module is reusable. Two enzyme catalysed reactions in series can be represented by the bond graph in Figure 8(d). This is equivalent to the reactions

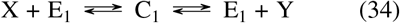

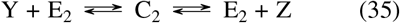

Larger networks of such two-state enzymes can be constructed through this approach. Furthermore, more complex enzyme-catalysed mechanisms could be implemented by replacing the **ECR** modules with different modules [14].

### 5.2. White-box modularity

For many applications in biology, it is beneficial for external ports to be defined in a flexible manner, prior to coupling models together. For example, a modeller may develop a model of a metabolic signalling pathway without *a priori* knowledge of the other pathways it may be connected to [62]. This allows individual models to be simulated and tested in isolation, while also being able to be merged with other models without connections being defined in advance.

Because bond graphs are graphical, they support this flexibility [14, 15, 17]. Any one-port component can be replaced by an external connection and then connected to an external module. For instance, the two enzyme reactions in Figure 8(d) could be represented separately as Figures 9(a) and (b). In a white-box modularity approach, the **Ce**:**Y** components are recognised as being equivalent, and can be merged through the following steps:

1. Disconnect the **Ce**:**Y** components in each module and replace them with **SS**:**Y** components.
2. Add the two enzyme modules to another bond graph, which also contains a **Ce**:**Y** component connected to a **0** junction. This ensures a common chemical potential as well as mass conservation.
3. Connect the two modules to the enzyme modules through the [Y] port.

**Figure 9:**
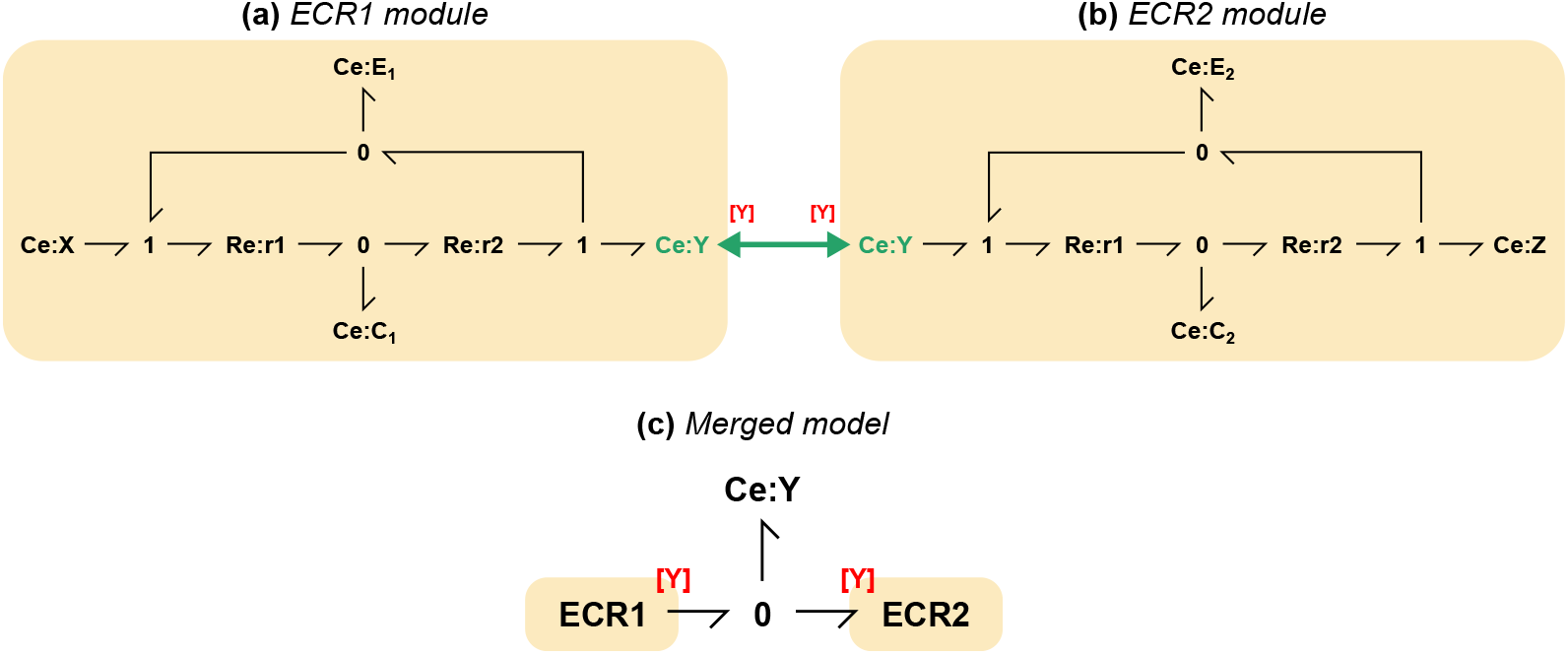
A white-box approach to model composition. (a) An enzyme catalysed reaction 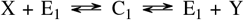, embedded within the **ECR1** module. (b) An enzyme catalysed reaction 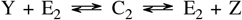, embedded within the **ECR2** module. During model merging, the **Ce**:**Y** component (shown in green) is common to both modules. It is therefore recognised as a point of connection. The components are then replaced by ports (indicated by the red [Y] labels). (c) The merged model connects the [Y] ports to a single **Ce**:**Y** component through a **0** junction to ensure mass conservation.

The resulting bond graph gives the same model as in Figure 9(c).

Model merging can be facilitated by unambiguously annotating species and reactions using ontological terms from databases [63]. Using this semantics-based approach, bond graph models can be systematically merged by recognising equivalent species and defining a set of graph-based rules to merge the individual bond graphs [17].

## 6. Stoichiometry and Bond Graphs

Stoichiometric analysis of biomolecular systems [64–67] looks at the null spaces of the *stoichiometric matrix* to derive fundamental properties of the systems expressed as *conserved moieties* and *flux paths*. Stoichiometric analysis is closely linked with the bond graph approach [2, 18].

The stoichiometric matrix be directly obtained from a bond graph [2, 3]; this is examined in § 6.1. Conversely, a bond graph can be obtained from the stoichiometric matrix [15]. Open systems can be created from closed systems using the concept of *chemostats* [4, 6, 68]; this is examined in § 6.2. The stoichiometric concept of pathways has a bond graph interpretation [6]; this is examined in § 6.3. In parallel with the seminal work of Oster et al. [35, 36], the mathematical foundations of chemical reaction networks (CRN) were being laid by Feinberg [69], Horn and Jackson [70] and Feinberg and Horn [71]; these results are collated by Feinberg [72]. This approach to chemical reaction network theory was further developed by Sontag [73], Angeli [74], and van der Schaft et al. [75, 76, 77]. The relation between CRNs and bond graphs is discussed by Gawthrop and Crampin [7]; this is examined in §6.4.

### 6.1. The stoichiometric matrix

In bond graph terms, a chemical reaction network, such as that depicted in Figure 4(a), can be thought of a number of reactions (represented by **Re** components) connected to species (represented by **Ce** components) by bonds. In particular, the network of bonds relates the reaction flows *f* to the the species flows 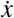. Collecting the *n_F_* reaction flows into a column vector *F* and the *n_X_* species flows into the column vector *x, x* can be expressed in terms of *F* as

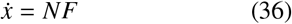

*N* is called the *stoichiometric matrix* and contains integer elements. For example, the stoichiometric matrix of the example in Figure 7 is

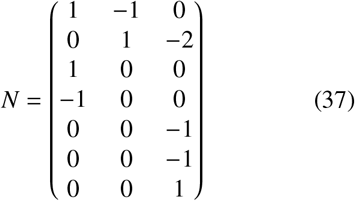

where

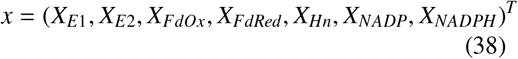

and *F* = (*v*_1_, *v*_2_, *v*_3_)^*T*^.

Equation (36) relates the reaction flows *F* to the species flows 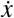. As the bond graph approach is energy based, the stoichiometric matrix N appearing in (36) can also be used to relate the corresponding energy covariables: the species chemical potentials φ and the reaction potentials Φ. In a closed system, the net power flowing into the **Ce** and **Re** components must be zero hence:

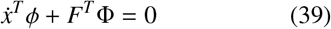

Using Equation (36), Equation (39) can be rewritten as:

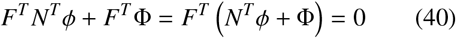

It follows that:

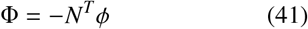

Thus not only does the stoichiometric matrix *N* relate the reaction flows *F* to the species flows 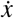, but its negative transpose relates the species chemical potentials *φ* and the reaction potentials Φ.

As discussed in § 3, it is necessary to decompose reaction potential Φ into the forward potential Φ^*f*^, and the reverse potential Φ^*r*^ where:

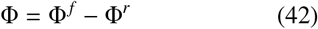

Defining *v^f^* and *v^r^* as the flow covariables of Φ^*f*^ and Φ^*r*^, *v^f^* and *v^r^* are the reaction flows corresponding the the left and right hand sides of the reaction equations. These are, of course, equal

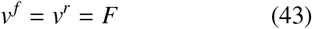

Nevertheless, it is convenient to decompose Equation (36) as:

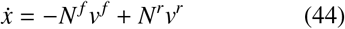

where *N^f^* and *N^r^* are the forward and reverse stoichiometric matrices respectively, containing only positive integer elements. Combining Equation (36), Equation (43) and Equation (44), it follows that

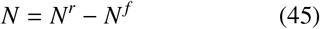

Also, using the same argument as that leading to Equation (41),

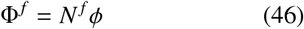

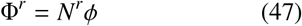

As discussed by Cellier and Greifeneder [40], the pair of equations (36) and (41) are equivalent to an (energy-transmitting) multiport transformer with modulus *N* – the stoichiometric matrix. This leads to the conceptual bond graph of Figure 10(a), where 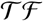 is used to represent a multiport transformer, 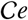 the **Ce** components and 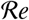 the **Re** components.

**Figure 10:**
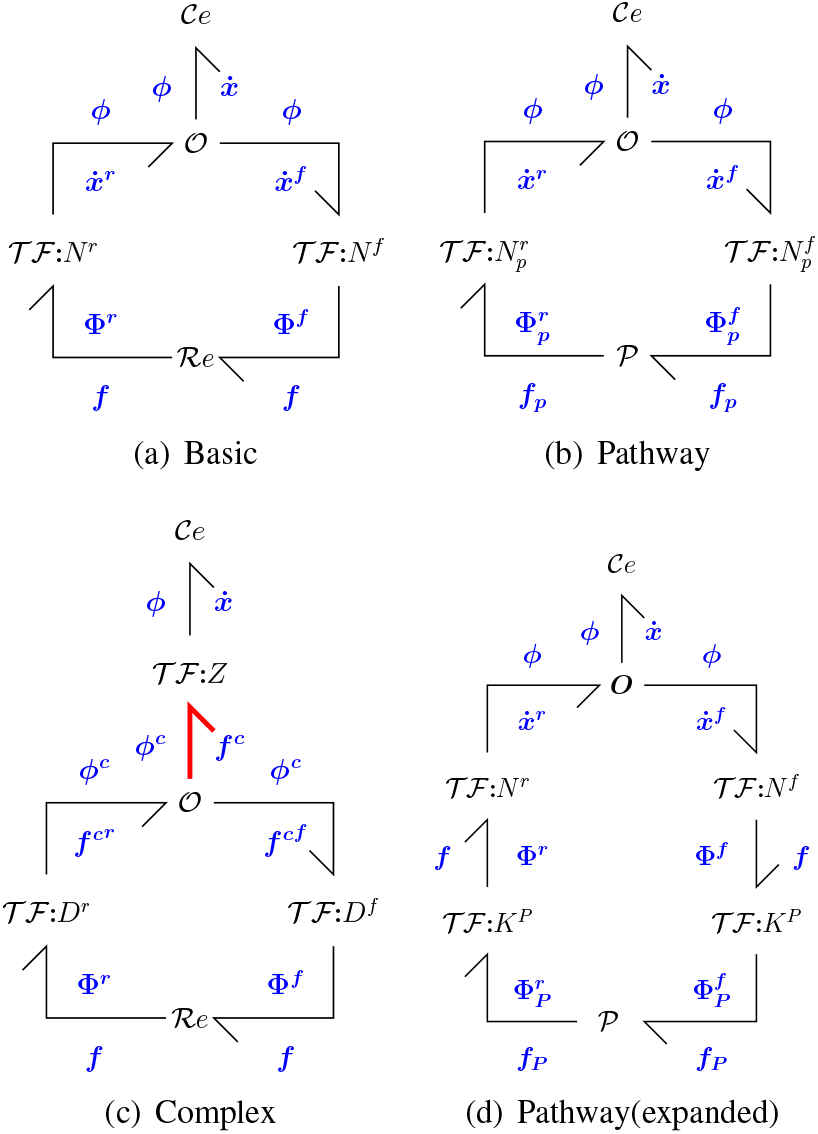
Stoichiometry of Bond Graphs

### 6.2. Chemostats and Open Systems

As discussed by Gawthrop and Crampin [4, 6], open biomolecular systems can be described and analysed using the notion of *chemostats* [4, 68]. Chemostats have two biomolecular interpretations:

1. one or more species are fixed to give a constant concentration (for example under a specific experimental protocol); this implies that an appropriate external flow is applied to balance the internal flow of the species.
2. as a **Ce** component with a fixed state imposed on a model in order to analyse its properties [3].

When chemostats are present, the reaction flows are determined by the dynamic part of the stoichiometric matrix. In this case the stoichiometric matrix *N* can be decomposed as the sum of two matrices [4]: the *chemostatic* stoichiometric matrix *N^cs^* and the *chemodynamic* stoichiometric matrix *N^cd^*

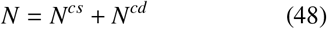

*N^cd^* is the same as *N* except that the *rows* corresponding to the chemostat variables are set to zero [6].

With these definitions, an open system can be expressed as

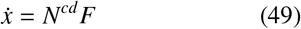

The stoichiometric properties of *N^cd^*, rather than *N*, determine system properties when chemostats are present.

### 6.3. Pathways

As discussed in the textbooks [64, 66, 67, 78], the (non-unique [6]) *n_F_* × *n_P_* null-space matrix *K_p_* of the open system stoichiometric matrix *N^cd^* (48) has the property that

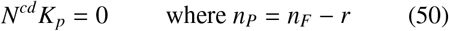

and *r* is the rank of *N. N^cd^* is an integer matrix and therefore *K_p_* can also be chosen to be an integer matrix. This is achieved by performing row reduction, as implemented in the `Matrix.nullspace()` method from the Python sympy package. Furthermore, if the reaction flows *F* are constrained in terms of the *n_P_* pathway flows *F_p_* as

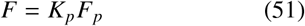

then substituting Equation (51) into Equation (36) and using Equation (50) implies that 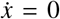. This is significant because the biomolecular system of equation (36) may be in a steady state for any choice of *F_p_*.

Following the arguments of § 6.1, Equation (51) can be interpreted as a bond graph multi-port transformer transmiting the energy flow 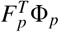 to *F^T^* Φ and thus represented by the conceptual bond graph of Figure 10(d). Defining

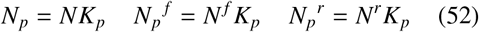

give the conceptual bond graph of Figure 10(b).

Continuing the example of Figure 7 with the reactions (30)–(32) and the chemostats are FdOx, FdRed, Hn, NADP and NADPH (in that order), the pathway matrix *K_p_* is

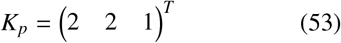

From Equation (52)

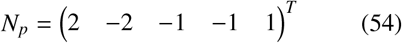

This gives the pathway reaction:

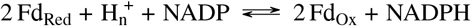

### 6.4. Chemical Complexes

The formal concept of *complexes* is essential to chemical reaction network theory [72]. Complexes are the combination of chemical species forming the substrate and products of the network reactions. This section summarises the links between chemical reaction network theory to the bond graph approach by incorporating the concept of complexes – further details are given by [7].

The complexes form the left and right hand sides of chemical reactions and each is associated with the coresponding reaction flow. Defining *Z* as the matrix relating the species flows 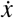 and the complex flows *f^c^*, 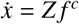. Further, defining *D* as the matrix relating the complex flows *f^c^* and the reaction flows *f, f^c^* = *Df*. Thus, 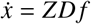 and the stoichiometric matrix can be decomposed as *N* = *ZD*. As in § 6.1, the matrices *Z* and *D* can be regarded as bond graph junctions and bonds represented as *transformers*; this is indicated in Figure 10(c).

As an example, consider the reactions (30)–(32) corresponding to the reaction network of Figure 7. The complexes corresponding to the three sets of reactants are: Fd_Red_, E_1_ and 2 E_2_ + Hn + NADP; the complexes corresponding to the three sets of products are: E_1_ + FdOx, E_2_ and NADPH. This gives six complexes in total^1^. The relations between the three reactions, the six complexes and the seven species are given graphically in Figure 11. The *Z* and *D* matrices are given in the Additional Material.

**Figure 11:**
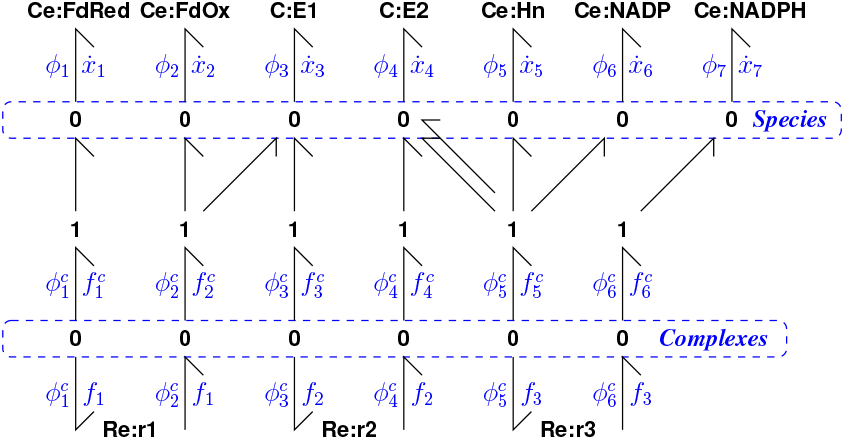
Ferredoxin reductase (Figure 7) in CRN form. The 7 species are mapped to the 6 complexes (Fd_red_, Fd_ox_ + E_1_, E_1_, E_2_, 2 E_2_ + H_n_ + NADP, NADPH) by the bonds connecting the 7 upper **0** junctions to the 6 lower **0** junctions; this mapping corresponds to the matrix Z. The 6 complexes are mapped onto the three reactions by the bonds connecting the lower **0** junctions to the **Re** components; this mapping corresponds to the matrix *D.*

## 7. Example: Photosynthesis

Photosynthesis within plant chloroplasts is the basis of life on earth [44–47]. As shown in Figure 12(a), the chloroplast has a membrane separating an inner space (lumen) from an outer space (stroma). In the chloroplast, the lumen gains protons and is called the p-space, the stroma loses protons and is called the n-space [45]. Thus, geometrically, the lumen corresponds to the mitochondrial matrix and the stroma to the mitochondrial inter-membrane space. However, electrically, the p-space is inside and the n-space outside – the reverse of the mitochondrial situation.

**Figure 12:**
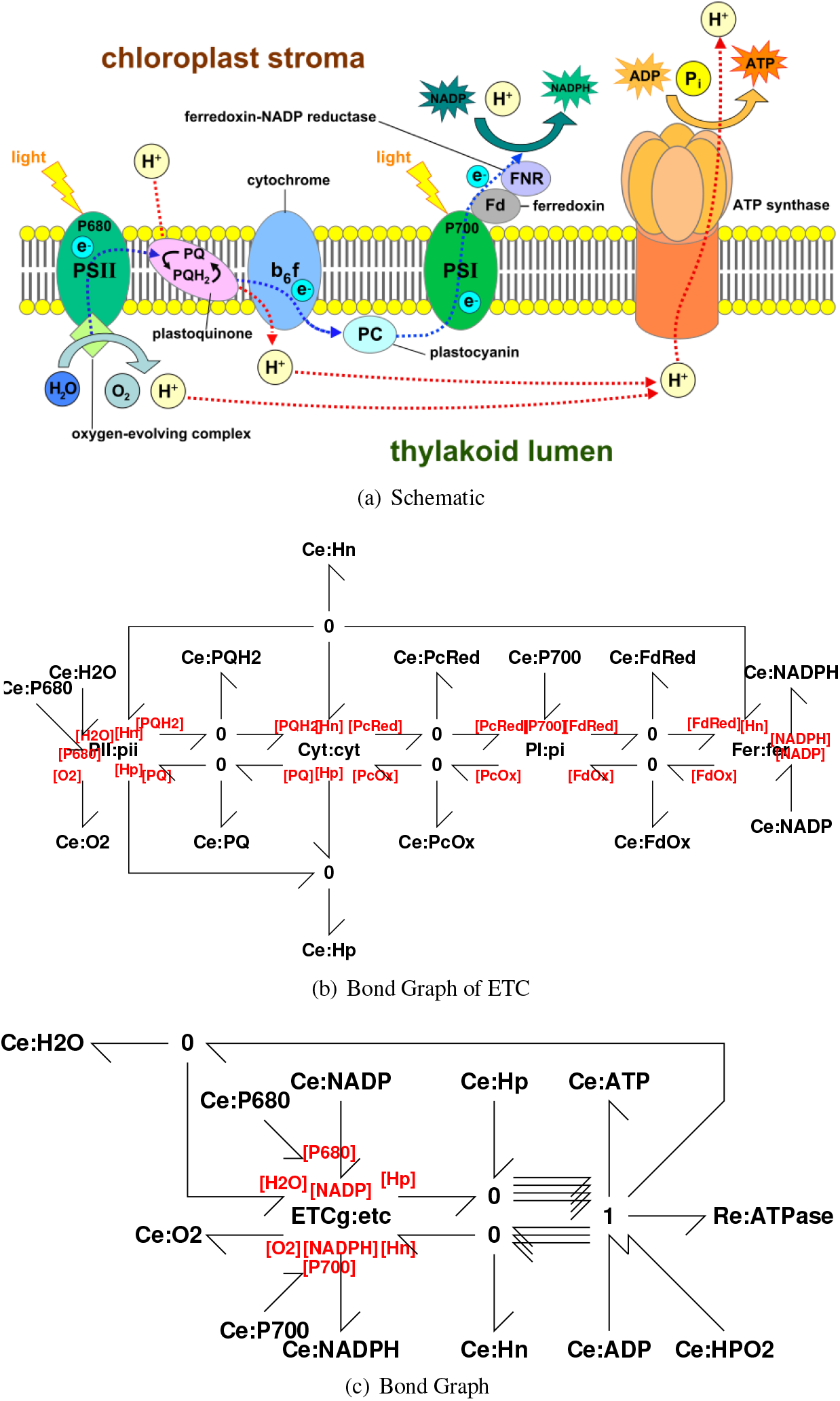
Photosynthesis. (a) Schematic representation (Created by Somepics and used under CC-BY-SA 4.0 https://commons.wikimedia.org/wiki/File:Thylakoid_membrane_3.svg). (b) Electron transport chain bond graph. The four modules Photosystem II (**PII**), Cytochrome (**CYT**), Photosystem I (**PI**) and Ferredoxin-NADP reductase (**Fer**) are described in the text; the bond graph representation for (**Fer**) is given in § 4.2, Figure 7 and the rest are given in the Additional Material. (c) Photosynthesis: generating NADPH, ATP and O_2_ from photons. The module **ETCg** conprises the Electron Tranport Chain of (b). Four lumen protons H_p_ produce one ATP from ADP and Pi consuming one stroma proton H_n_ and producing one H_2_O; the remaining three stroma H_n_ protons are returned.

### 7.1. Modular bond graph model

The chloroplast electron transport chain (ETC) has 4 complexes, each of which is represented by the bond graph modules of Figure 12(b): Photosystem II (**PII**), Cytochrome (**CYT**), Photosystem I (**PI**) and Ferredoxin - NADP reductase (**Fer**). As an example, the bond graph for **Fer** is given in § 4.2, Figure 7; the bond graphs for each of the other three modules is given in the Additional Material. The four modules were coupled using the white-box approach described in § 5.2.

**PII** Photosystem II absorbs photons (P_**680**_) at wavelength 680nm and splits water, releasing protons into the p-space and passing electrons to the plastoquinone (PQ) – plastoquine (PQH2) couple which absorbs protons from the n-space. The net reaction is:

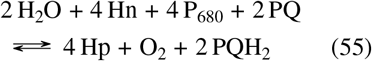 A more detailed model would include mechanisms of photon absorption [44].
**Cyt** Cytochrome passes electrons to the plastoquine – plastoquinone couple which releases two protons into the p-space. Electrons are passed to the plastocyanine couple (PcOx – PcRed). Two protons are pumped across the membrane. The net reaction is:

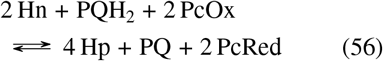
**PI** Photosystem I absorbs photons (P_700_) at wavelength 700nm and transports electrons from the plastocyanine (PcRed – PcOx) couple to the ferredoxin (FdOx – FdRed) couple. The net reaction is:

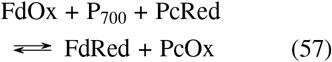
**Fer** Ferredoxin-NADP reductase transfers electrons from the ferredoxin (FdRed – FdOx) couple to convert NADP to NADPH absorbing a proton from the n-space. The net reaction is:

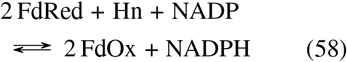

Applying the pathway analysis of § 6.3, the modular Electron transport chain has the overall reaction:

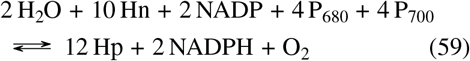

As well as generating NADPH from NADP and splitting water, the reaction passes 12 protons from the negative to the positive space. These are used to generate ATP as shown in Figure 12(c), where four protons are required to generate each ATP molecule [46]. Applying the pathway analysis of § 6.3, the modular bond graph of Figure 12(c) has the overall reaction:

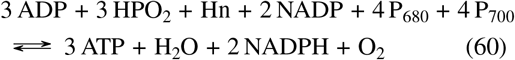

This reaction corresponds to the statement: “The absorption of eight photons yields one O_2_, two NADPH and three ATP molecules” [46, § 19.4].

### 7.2. Energetics

Photosynthesis, as summarised by reaction (60), can be thought of as converting the energy of photons P_680_&P_700_ to the chemical energy of products such as NADPH and ADP. By estimating the input photon energy and the net energy of products, it is possible to obtain an estimate of conversion efficiency.

The standard formula [44] for the energy (J) of a photon is

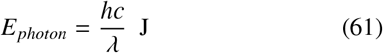

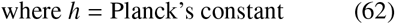

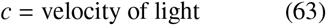

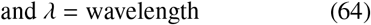

This can be converted into energy per coulomb (J C^-1^ or V) of photons to give

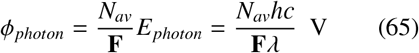

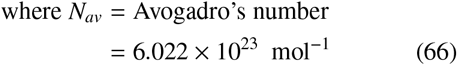

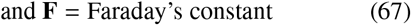

For example:

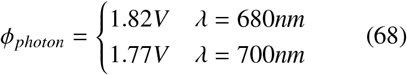

Thus, from Reaction (60), the energy input corresponding to one O_2_ molecule is

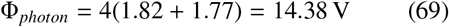

**Table 2:**
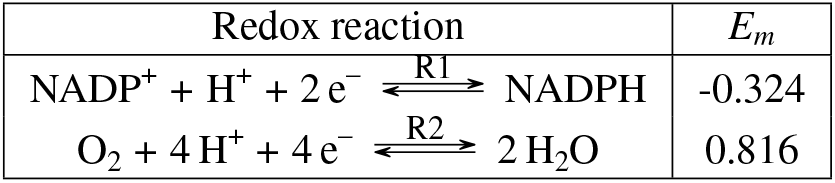
Redox potentials (at pH 7) from [44, Table A1.2].

A key challenge in the development of dynamic models is the fitting of parameters to experimental data, especially when thermodynamic constraints need to be satisfied. Here, we use the thermodynamically safe parameterisation provided by bond graphs to resolve this issue [15]. Parameter estimation depends on both the form of the model and the type of data available. For example, a bond graph model combined with reaction potential Φ data can be used to estimate a consistent set of species potentials *ϕ* [15]. Moreover, reaction potentials provide a route to computing the *efficiency* of a chain of reactions [9]. Using the close relationship between redox potentials and reaction potential [19], this section combines the bond graph model of this paper with redox data for photosynthesis [44]. In particular, for a given half-reaction, the reaction potential Φ is given in terms of the redox potential *E_m_* by

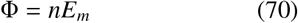

where n is the number of electrons. Thus from Table 2, the potentials for the reactions R1 and R2 are:

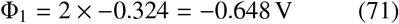

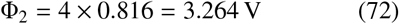

ATP hydrolysis (in the stroma) has the reaction R3:

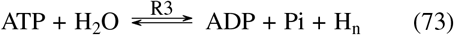

Thus the *chemical* part of the chloroplast reaction (60) can be decomposed in terms of the two reactions of Table 2 and reaction (73) as:

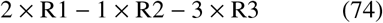

According to [59, Case study 4.2], the standard (at pH 7) Gibbs energy of reaction R3 is A_3_ = 34 kJ mol^-1^. From § 4.1 it follows that:

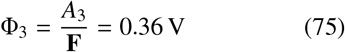

Hence the overall potential of the chemical products of the reaction is the negative of the potential of the chemical part of the reaction:

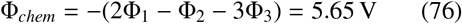

Following [9], the efficiency *η* can be defined as the ratio of the potential of the products to input driving potential:

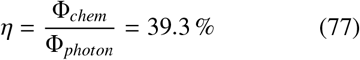

The reaction potentials all correspond to standard conditions; thus changing the concentarations would change the product potential Φ_*chem*_ and thus the efficiency *η*. There are many possible definitions of photosynthesis efficiency; one of these is *Energy Storage Efficiency* which is given by [44, § 13.3] as ≈ 27 %. As this is based on the final carbohydrate products this would be expected to be lower than that of (77).

## 8. Future research directions

The work of Edmund Crampin’s group summarised in this paper provides the basis for energy based modelling of living systems at all scales from the cell to the human physiome. Such multi-scale integrated models are have great potential for science, medicine and biotechnology. To achieve this promise, there is a need for research in a number of areas.

The large amount of high-throughput data now available, such as proteomics, metabolomics, lipidomics and fluxomics, could be used to estimate the numerical parameters for energy based models. While initial work in this area has made use of fluxomics data [15], further work is required to incorporate other other biological data and to account for discrepancies between datasets. To help characterise uncertainty, sensitivity analysis of bond graph models [79] needs to be extended to large biological systems. Parameter uncertainty in the probabilistic sense, such as that employed in parameter balancing [51], is also expected to play a role here.

Living systems contain many feedback loops. These are essential to physiological function, and form the basis for constructing networks in synthetic biology. However, in contrast to traditional control systems, biological controllers are themselves physical components and are subject to fundamental limitations in how they operate. Initial work has characterised the use of passive control in biology, as opposed to active control in engineering [23]. Future work will extend this work to integrate control-theoretical concepts and apply them in the context of synthetic biology. Some synthetic constructs have been observed to compete with the endogenous pathways within the cell for resources [80]. An energy-based approach has the potential to optimise the performance of synthetic circuits while minimally affecting the natural pathways inside a cell.

Living systems have a significant spatial dimension including cell motility, blood flow and heterogenous organs [81]. The port-Hamiltonian is a spatial extension of the bond graph which has recently been applied to biology [82]. However, modelling biochemistry in this context remains the subject of further work. Once the extension to spatial domains has been achieved, we will be able to combine high-throughput omics data with imaging data to develop detailed, cell-specific models for medicine and biotechnology [83].

## Supporting information

Supplementary material part 1

Supplementary material part 2

## Code availability

The code for this paper is available at https://github.com/gawthrop/GawPan22.

## Footnotes

1 In general the complexes may not all be different [7].

