## Supplementary material part 1 for "Network Thermodynamics of Biological Systems: A Bond Graph Approach"

### Additional material for:
### Network Thermodynamics of Biological Systems: A Bond Graph Approach.
### *Part 1: Reaction modelling*

Peter Gawthrop and Michael Pan

May 4, 2022

### 1 Introduction

- This document contains additional material for the paper: *Network Thermodynamics of Biological Systems: A Bond Graph Approach* by Peter Gawthrop and Michael Pan.
- It illustrates how the Python package BondGraphTools can be used to create and analyse chemical reaction networks.
- It provides background to Section 3, Chemical Reactions.
- This document is **Reactions.pdf** (see also **Chloroplast.pdf**).
- The document is available as the Jupyter notebook **Reactions.ipynb**.
- Code is available at `https://github.com/gawthrop/GawPan22`

#### 1.1 Some useful imports

[1]:
```
## Some useful imports
import BondGraphTools as bgt
import numpy as np
import sympy as sp
import matplotlib.pyplot as plt

## Stoichiometric analysis
import stoich as st

## SVG bg representation conversion
import svgBondGraph as sbg

## Modular bond graphs
import modularBondGraph as mbg

## Display (eg disp.SVG(), disp.
import IPython.display as disp

## Data
import phiData
import redoxData

import importlib as imp

## Some switches
chemformula = True
quiet = True
```

### 2 Basic reactions

## 2.1 A ⇔ B

##### 2.1.1 Graphical bond graph representation

[2]:
```
disp.SVG('ABbasic_abg.svg')
```

[2]:

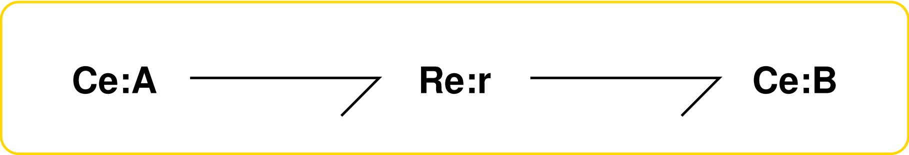

##### 2.1.2 Convert graphical representation to BondGraphTools representation

[3]:
```
# imp.reload(sbg)
sbg.model('ABbasic_abg.svg',quiet=quiet)
import ABbasic_abg
# imp.reload(ABbasic_abg)
```

##### 2.1.3 Stoichiometric analysis and reactions

[4]:
```
## Stoichiometry
sABbasic = st.stoich(ABbasic_abg.model(),quiet=quiet)
disp.Latex(st.sprintrl(sABbasic,chemformula=chemformula))
```

[4]:

$$\mathrm{A} \overset{r}{\rightleftharpoons} \mathrm{B} \tag{1}$$

### 3 Reactions with connections

## 3.1 A + B ⇔ C + D

##### 3.1.1 Graphical bond graph representation

[5]:
```
disp.SVG('ABCDbasic_abg.svg')
```

[5]:

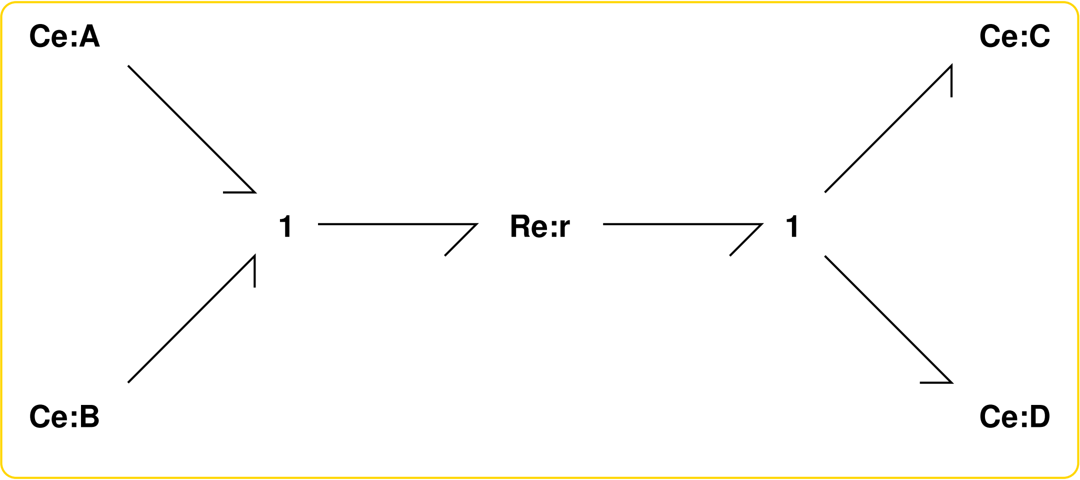

##### 3.1.2 Convert graphical representation to BondGraphTools representation

[6]:
```
# imp.reload(sbg)
sbg.model('ABCDbasic_abg.svg',quiet=quiet)
import ABCDbasic_abg
# imp.reload(ABCDbasic_abg)
```

##### 3.1.3 Stoichiometric analysis and reactions

[7]:
```
## Stoichiometry
sABCDbasic = st.stoich(ABCDbasic_abg.model(),quiet=quiet)
disp.Latex(st.sprintrl(sABCDbasic,chemformula=chemformula))
```

[7]:

$$\mathtt{A} + \mathtt{B} \xrightleftharpoons{\mathtt{r}} \mathtt{C} + \mathtt{D} \tag{2}$$

## 3.2 A ⇔ B ⇔ C

##### 3.2.1 Graphical bond graph representation

[8]:
```
disp.SVG('ABCbasic_abg.svg')
```

[8]:

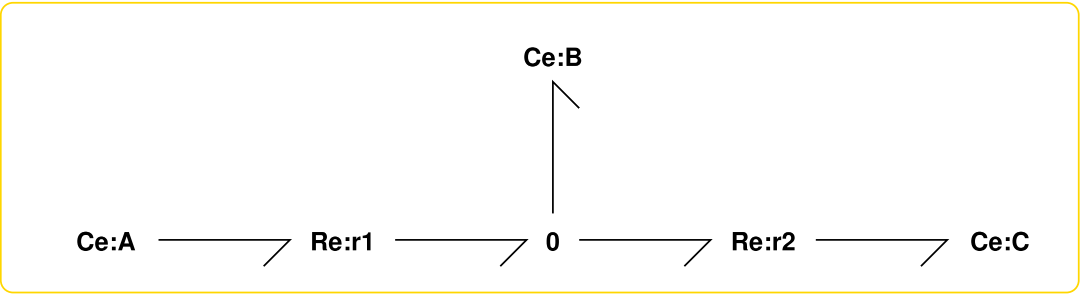

##### 3.2.2 Convert graphical representation to BondGraphTools representation

[9]:
```
# imp.reload(sbg)
sbg.model('ABCbasic_abg.svg',quiet=quiet)
import ABCbasic_abg
# imp.reload(ABCbasic_abg)
```

##### 3.2.3 Stoichiometric analysis and reactions

[10]:
```
## Stoichiometry
sABCbasic = st.stoich(ABCbasic_abg.model(),quiet=quiet)
disp.Latex(st.sprintrl(sABCbasic,chemformula=chemformula))
```

[10]:

$$\mathtt{A} \underset{}{\overset{\mathtt{r}_1}{\rightleftharpoons}} \mathtt{B} \tag{3}$$

$$\mathtt{B} \underset{}{\overset{\mathtt{r}_2}{\rightleftharpoons}} \mathtt{C} \tag{4}$$

### 4 Multiple stoichiometry

## 4.1 A+2B ⇔ 3C+4D

##### 4.1.1 Graphical bond graph representation

[11]:
```
disp.SVG('ABCDmult_abg.svg')
```

[11]:

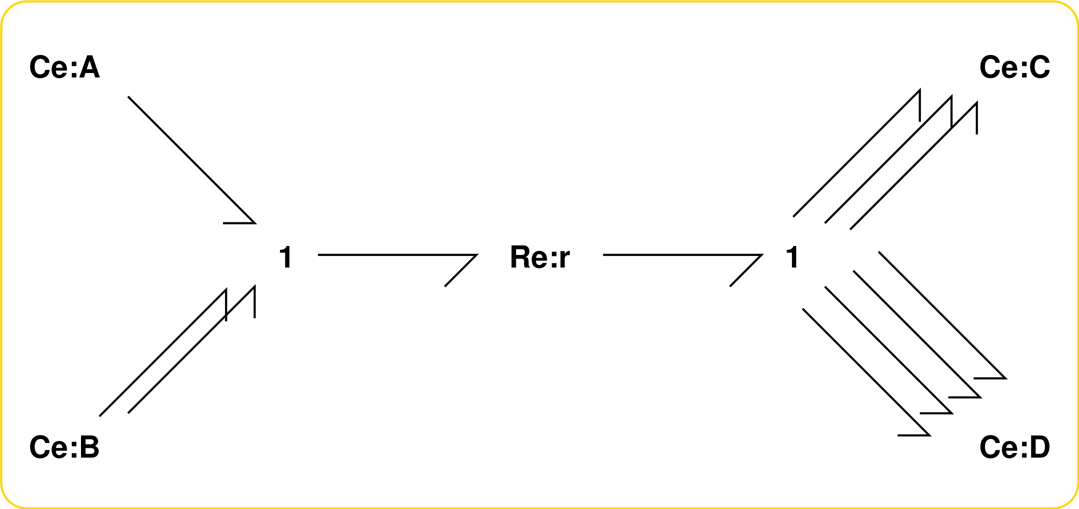

##### 4.1.2 Convert graphical representation to BondGraphTools representation

[12]:
```
# imp.reload(sbg)
sbg.model('ABCDmult_abg.svg',convertCe=True,quiet=quiet)
import ABCDmult_abg
# imp.reload(ABCDmult_abg)
```

##### 4.1.3 Stoichiometric analysis and reactions

[13]:
```
## Stoichiometry
sABCDmult = st.stoich(ABCDmult_abg.model(),quiet=quiet)
disp.Latex(st.sprintrl(sABCDmult,chemformula=chemformula))
```

[13]:

$$\mathrm{A} + 2\,\mathrm{B} \overset{r}{\rightleftharpoons} 3\,\mathrm{C} + 4\,\mathrm{D} \tag{5}$$

### 5 One-port Re component

## 5.1 A+2B ⇔ 3C+4D

##### 5.1.1 Graphical bond graph representation

[14]:
```
disp.SVG('ABCDone_abg.svg')
```

[14]:

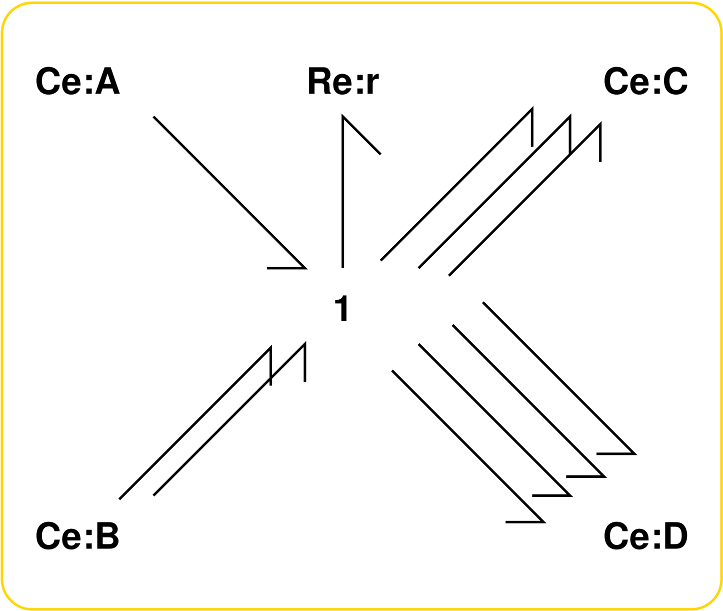

##### 5.1.2 Convert graphical representation to BondGraphTools representation

[15]:
```
# imp.reload(sbg)
sbg.model('ABCDone_abg.svg',convertCe=True,convertR=True,quiet=quiet)
import ABCDone_abg
# imp.reload(ABCDone_abg)
```

##### 5.1.3 Stoichiometric analysis and reactions

[16]:
```
## Stoichiometry
sABCDone = st.stoich(ABCDone_abg.model(),quiet=quiet)
disp.Latex(st.sprintrl(sABCDone,chemformula=chemformula))
```

[16]:

$$\mathtt{A + 2\,B \xrightleftharpoons{r} 3\,C + 4\,D} \tag{6}$$
