## Supplementary material part 2 for "Network Thermodynamics of Biological Systems: A Bond Graph Approach"

### Additional material for: Network Thermodynamics of Biological Systems: A Bond Graph Approach

*Part 2: Photosynthesis modelling.*

May 4, 2022

### 1 Introduction

- This document contains additional material for the paper: *Network Thermodynamics of Biological Systems: A Bond Graph Approach* by Peter Gawthrop and Michael Pan.
- It illustrates how the Python package BondGraphTools can be used to create and analyse chemical reaction networks.
- It provides background to Section 4.2 *Redox Reactions*, Section 6 *Stoichiometry and Bond Graphs* and Section 7 *Example: Photosynthesis*.
- This document is **Chloroplast.pdf** (see also **Reactions.pdf**).
- The document is available as the Jupyter notebook **Chloroplast.ipynb**.
- Code is available at `https://github.com/gawthrop/GawPan22`

#### 1.1 Some useful imports

[1]:
```
## Some useful imports
import BondGraphTools as bgt
print('Using BondGraphTools', bgt.version)
import numpy as np
import sympy as sp
import matplotlib.pyplot as plt

## Stoichiometric analysis
import stoich as st

## SVG bg representation conversion
import svgBondGraph as sbg

## Stoich to BG
import stoichBondGraph as stbg

## Modular bond graphs
import modularBondGraph as mbg

## Display (eg disp.SVG(), disp.
import IPython.display as disp

## Data
import phiData
import redoxData

## Copy
import copy

import importlib as imp

quiet = True
chemformula = True
savefig = False
```

```
Using BondGraphTools 0.3.7
```

[2]:
```python
def SaveFig(savefig,name):
    if savefig:
        print('Saving',name)
        plt.savefig(name)
    else:
        print('Not saving',name)
```

[3]:
```python
def mV(E):
    return int(round(1000*E))
```

##### 1.2 Photosynthesis

Photosynthesis within plant chloroplasts is the basis of life on earth Blankenship (2021), Nicholls and Ferguson (2013).

Like the mitochondrion, the chloroplast has a membrane seperating an inner space (lumen) from an outer space (stroma). In the chloroplast, the lumen gains protons and is called the p-space, the stroma looses protons and is called the n-space. Thus geometrically, the lumen corresponds to the mitochondrial matrix and the stroma to the mitochondrial intermembrane space; but electrically the p-space is inside and the n-space outside - the reverse of the mitochondrial situation.

The chloroplast electron transport chain has 4 complexes.

1. Photosystem II (PII) which absorbs photons at 680nm and splits water releasing protons into the p-space and passing electrons to the plastoquinone(PQ)/plastoquine(PQH2) couple which absorbs protons from the n-space.
2. Cytochrome bf (Cyt) which passes electrons to the plastoquine/plastoquinone couple which releases two protons into the p-space. Electrons are passed to the plastocyanine couple (PcOx/PcRed). Two protons are pumped across the membrane.
3. Photosystem I (PI) which absorbs photons at 700nm and transports electrons from the plastocyanine (PcRed/PcOx) couple to the ferredoxin (FdOx/FdRed) couple.
4. Ferredoxin-NADP reductase which transfers electrons from the ferredoxin (FdRed/FdOx) couple to convert NADP to NADPH absorbing a proton from the n-space.

The following figure is: https://commons.wikimedia.org/wiki/File:Thylakoid_membrane_3.svg

- See section 7 of paper.

[4]:
```python
# https://commons.wikimedia.org/wiki/File:Thylakoid_membrane_3.svg
disp.SVG("Thylakoid_membrane_3.svg")
```

[4]:

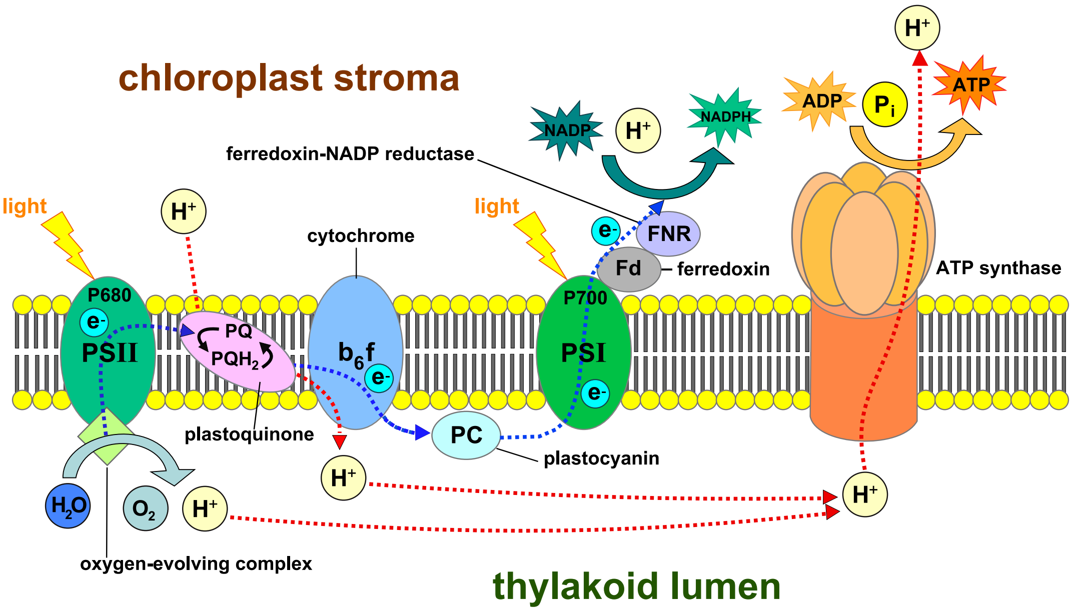

#### 2 Photon Energetics

- See section 7.2 of paper.

$$\phi_{photon} = \frac{N_{av}hc}{F\lambda} \tag{1}$$
$$\text{where } N_{av} = \text{Avogadro's number} \tag{2}$$
$$h = \text{Planck's constant} \tag{3}$$
$$c = \text{velocity of light} \tag{4}$$
$$F = \text{Faraday's constant} \tag{5}$$
$$\text{and } \lambda = \text{wavelength} \tag{6}$$

For example:

$$\phi_{photon} = \begin{cases} 1.82V & \lambda = 680nm \\ 1.77V & \lambda = 700nm \end{cases} \tag{7}$$

[5]:
```
## Plot
wavelength = np.linspace(600,800,100)
phi_680 = redoxData.V_photon(680)
phi_700 = redoxData.V_photon(700)
print(f'phi_680 = {phi_680:0.2f}')
print(f'phi_700 = {phi_700:0.2f}')
phi_wave = [phi_680,phi_700]
wave = [680,700]
one = np.ones(2)

PHI = redoxData.V_photon(wavelength)
plt.plot(wavelength,PHI)
```

```
plt.hlines(phi_wave,min(wavelength)*one,wave,linestyles='dotted')
plt.vlines(wave,min(PHI)*one,phi_wave,linestyles='dotted')
plt.grid()
plt.xlabel('$\lambda$(nm)')
plt.ylabel('$V_{photon}$ V')
SaveFig(savefig,'Figs/V_photon.pdf')
```

```
phi_680 = 1.82
phi_700 = 1.77
Not saving Figs/V_photon.pdf
```

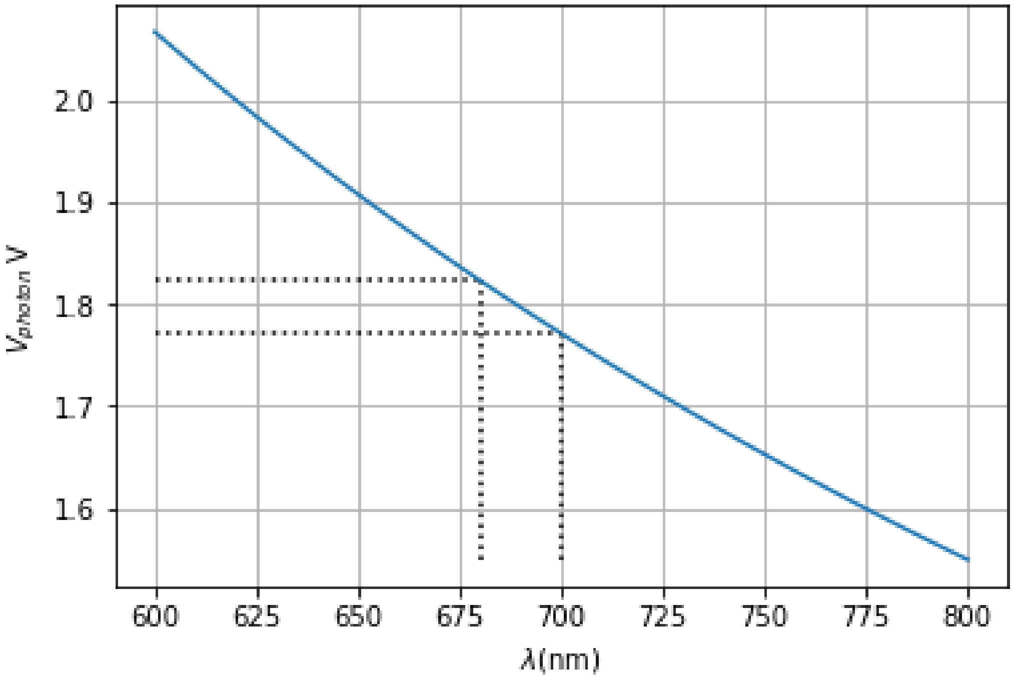

### 3 Redox reactions

#### 3.1 2 port Re version

- See section 3.5 of paper.

[6]:
```
## 2 port Re version
disp.SVG('Fer0_abg.svg')
```

[6]:

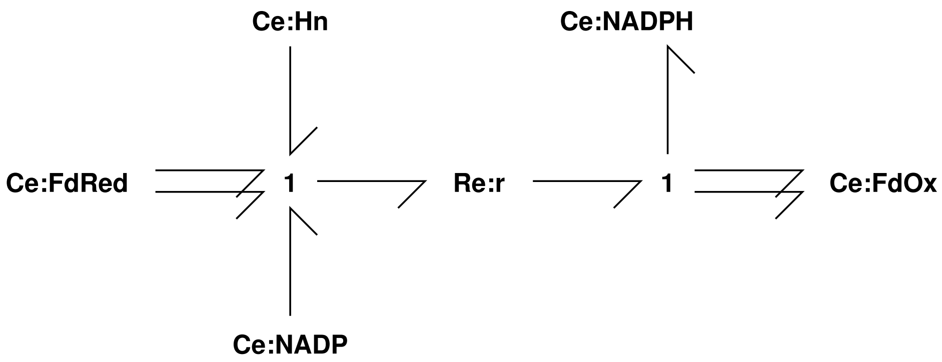

##### 3.2 1 port Re version

- See section 3.6 of paper.

[7]:
```
## One-port Re version
disp.SVG('Fer1_abg.svg')
```

[7]:

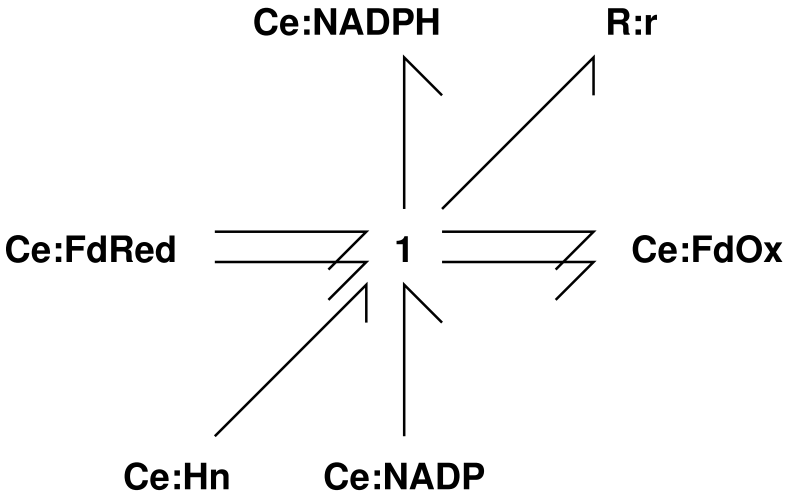

[8]:
```
## Convert to BGtools
sbg.model('Fer1_abg.svg',convertCe=True,convertR=True,quiet=quiet)
import Fer1_abg
```

[9]:
```
## Stoichiometry
sFer1= st.stoich(Fer1_abg.model(),quiet=True)
disp.Latex(st.sprintrl(sFer1,chemformula=chemformula))
```

[9]:

$$2\,\mathtt{FdRed} + \mathtt{Hn} + \mathtt{NADP} \xrightleftharpoons[]{\mathtt{r}} 2\,\mathtt{FdOx} + \mathtt{NADPH} \tag{8}$$

##### 3.3 Redox formulation

This *redox reaction* can be written as two half-reactions with the following BG representation:

[10]:
```
disp.SVG('Fer_abg.svg')
```

[10]:

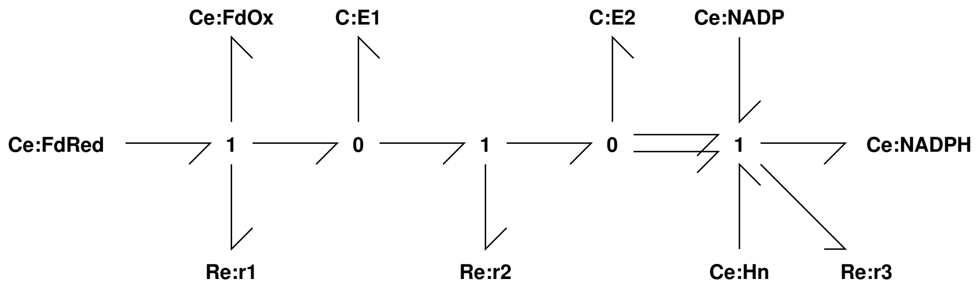

[11]:
```
sbg.model('Fer_abg.svg',convertCe=True,convertR=True,quiet=quiet)
import Fer_abg
# imp.reload(Fer_abg)
```

###### 3.3.1 Stoichiometry

[12]:
```
## Stoichiometry
sFer= st.stoich(Fer_abg.model(),quiet=True)
disp.Latex(st.sprintrl(sFer,chemformula=chemformula))
```

[12]:

$$\mathtt{FdRed} \xrightleftharpoons[]{\mathtt{r}_1} \mathtt{E}_1 + \mathtt{FdOx} \tag{9}$$

$$\mathtt{E}_1 \xrightleftharpoons[]{\mathtt{r}_2} \mathtt{E}_2 \tag{10}$$

$$2\,\mathtt{E}_2 + \mathtt{Hn} + \mathtt{NADP} \xrightleftharpoons[]{\mathtt{r}_3} \mathtt{NADPH} \tag{11}$$

###### 3.3.2 Pathway analysis

[13]:
```
## Pathway analysis
chemostats = ['FdRed','FdOx','NADP','NADPH','Hn']
scFer = st.stoich(Fer_abg.model(),chemostats=chemostats,quiet=True)
spFer = st.path(sFer,scFer)
disp.Latex(st.sprintrl(spFer,chemformula=chemformula))
```

[13]:

$$2\,\mathtt{FdRed} + \mathtt{Hn} + \mathtt{NADP} \xrightleftharpoons[]{\mathtt{pr}_1} 2\,\mathtt{FdOx} + \mathtt{NADPH} \tag{12}$$

### 4 Chemical reaction network (CRN) representation

- See section 6.4 of paper.

[14]:
```
disp.Latex(st.sprintl(sFer,'species'))
```

[14]:

$$X = \begin{pmatrix} X_{E1} \\ X_{E2} \\ X_{FdOx} \\ X_{FdRed} \\ X_{Hn} \\ X_{NADP} \\ X_{NADPH} \end{pmatrix} \tag{13}$$

[15]:
```
disp.Latex(st.sprintl(sFer,'N'))
```

[15]:

$$N = \begin{pmatrix} 1 & -1 & 0 \\ 0 & 1 & -2 \\ 1 & 0 & 0 \\ -1 & 0 & 0 \\ 0 & 0 & -1 \\ 0 & 0 & -1 \\ 0 & 0 & 1 \end{pmatrix} \tag{14}$$

[16]:
```
disp.Latex(st.sprintl(sFer,'Z'))
```

[16]:

$$Z = \begin{pmatrix} 0 & 1 & 0 & 1 & 0 & 0 \\ 0 & 0 & 2 & 0 & 1 & 0 \\ 0 & 0 & 0 & 1 & 0 & 0 \\ 1 & 0 & 0 & 0 & 0 & 0 \\ 0 & 0 & 1 & 0 & 0 & 0 \\ 0 & 0 & 1 & 0 & 0 & 0 \\ 0 & 0 & 0 & 0 & 0 & 1 \end{pmatrix} \tag{15}$$

[17]:
```
disp.Latex(st.sprintl(sFer,'D'))
```

[17]:

$$D = \begin{pmatrix} -1 & 0 & 0 \\ 0 & -1 & 0 \\ 0 & 0 & -1 \\ 1 & 0 & 0 \\ 0 & 1 & 0 \\ 0 & 0 & 1 \end{pmatrix} \tag{16}$$

[18]:
```
disp.SVG('FerCRN_abg.svg')
```

[18]:

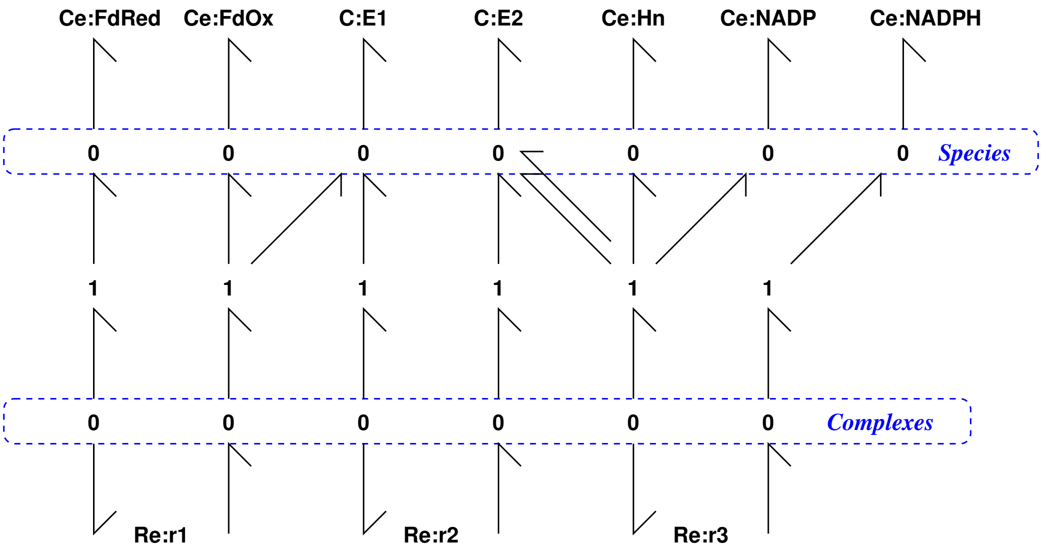

[19]:
```
sbg.model('FerCRN_abg.svg',quiet=quiet)
import FerCRN_abg
# imp.reload(FerCRN_abg)
```

#### 4.1 Stoichiometry

[20]:
```
## Stoichiometry
imp.reload(st)
sFerCRN= st.stoich(FerCRN_abg.model(),quiet=quiet)
disp.Latex(st.sprintrl(sFerCRN,chemformula=chemformula))
```

[20]:

$$\mathtt{FdRed} \xrightleftharpoons{r_1} \mathtt{E}_1 + \mathtt{FdOx} \tag{17}$$

$$\mathtt{E}_1 \xrightleftharpoons{r_2} \mathtt{E}_2 \tag{18}$$

$$2\,\mathtt{E}_2 + \mathtt{Hn} + \mathtt{NADP} \xrightleftharpoons{r_3} \mathtt{NADPH} \tag{19}$$

#### 4.2 Pathway analysis

[21]:
```
## Pathway analysis
chemostats = ['FdRed','FdOx','NADP','NADPH','Hn']
scFerCRN = st.stoich(FerCRN_abg.model(),chemostats=chemostats,quiet=True)
spFerCRN = st.path(sFerCRN,scFerCRN)
disp.Latex(st.sprintrl(spFerCRN,chemformula=chemformula))
```

[21]:

$$2\,\mathtt{FdRed} + \mathtt{Hn} + \mathtt{NADP} \xrightleftharpoons{pr_1} 2\,\mathtt{FdOx} + \mathtt{NADPH} \tag{20}$$

### 5 Redox potentials and equivalent species potentials $\phi$

Note that photon energies have been used for V_700 and V_680; this value is too large as the energy conversion is not direct. A model of this needs to be built. See, for example: Blankenship and Prince (1985)

According to Blankenship (2021) (sec. 8.3) the *electrical* part of the membrane potential is small in isolated choroplasts.

[22]:
```
#disp.Latex(redoxData.table())
```

[23]:
```
import redoxData
## pH from BerTymStr 19.3
pH_p = 4 # pH of p-space
pH_n = pH_p + 3
VpH = redoxData.VpH(pH_p - pH_n)
V680 = redoxData.V_photon(wavelength=680)
V700 = redoxData.V_photon(wavelength=700)
print(VpH)
print('V_pH =',int(1000*VpH),'mV')
print('V_680 =',int(1000*V680),'mV')
print('V_700 =',int(1000*V700),'mV')
```

```
0.18462122057715774
V_pH = 184 mV
V_680 = 1823 mV
V_700 = 1771 mV
```

[24]:
```
## Convert redox potentials to species phi
phi_redox = redoxData.phi()
phi_redox['Hn'] = redoxData.VpH(pH_n)
phi_redox['Hp'] = redoxData.VpH(pH_p)

## The electrical membrane potential is close to zero ()
phi_redox['dV'] = VpH

## Put in my computed values for photon potentials
phi_redox['P680'] = phi_680
phi_redox['P700'] = phi_700
```

[25]:
```
## phi for ATP etc from Phi for hydrolysis
# Phi_Hyd = 0.424
Phi_Hyd = 35e3/st.F()
print(f'Phi_hyd = {Phi_Hyd:0.2f}')
phi_H2O = phi_redox['H2O']
phi_H = phi_redox['Hn']
Phi_A = Phi_Hyd - phi_H2O + phi_H
print(f'Phi_A: {Phi_A:0.2f}')

phi_ATP = Phi_A/2
phi_ADP = phi_Pi = -Phi_A/4
```

```python
phi_redox['ATP'] = phi_ATP
phi_redox['ADP'] = phi_ADP
phi_redox['Pi'] = phi_Pi

## Sanity check
Phi_Hyd_check = phi_ATP + phi_H2O -(phi_ADP + phi_Pi + phi_H)
print(f'{Phi_Hyd_check-Phi_Hyd:0.2f}')
```

```
Phi_hyd = 0.36
Phi_A: 1.18
0.00
```

[26]:
```python
# Print values
for spec in phi_redox.keys():
    if not (('*' in spec) or ('E' in spec) or ('+' in spec)):
        print(f'phi_{spec} = {phi_redox[spec]:0.2f} V')
```

```
phi_NADP = -0.11 V
phi_NADPH = 0.11 V
phi_NAD = -0.10 V
phi_NADH = 0.10 V
phi_O2 = 2.49 V
phi_H2O = -1.25 V
phi_P700 = 1.77 V
phi_P870 = 0.45 V
phi_P680 = 1.82 V
phi_PQ = 0.43 V
phi_PQH2 = -0.43 V
phi_PcOx = 0.19 V
phi_PcRed = -0.19 V
phi_FdOx = -0.21 V
phi_FdRed = 0.21 V
phi_Hn = -0.43 V
phi_Hp = -0.25 V
phi_dV = 0.18 V
phi_ATP = 0.59 V
phi_ADP = -0.29 V
phi_Pi = -0.29 V
```

[27]:
```python
## Compatibility with Chloroplast model (but makes no difference)
phi_redox['etc_FdRed'] = phi_redox['FdRed']
phi_redox['etc_PcRed'] = phi_redox['PcRed']
phi_redox['etc_PQH2'] = phi_redox['PQH2']
```

[28]:
```python
## Redox potentials with zero photon contribution
phi_redox_0 = copy.copy(phi_redox)
phi_redox_0['P680'] = phi_redox_0['P700'] = 0
```

### 6 Bond graph description of ETC complexes

- The four ETC complexes are represented by SVG graphics which are automatically converted into BondGraphTools format.
- The stoichiometric toolbox is then used to generate the pathway-reduced equation for the complex.
- See section 7 of paper.

#### 6.1 Complex PII – Photosystem II

[29]:
```
disp.SVG('PII_abg.svg')
```

[29]:

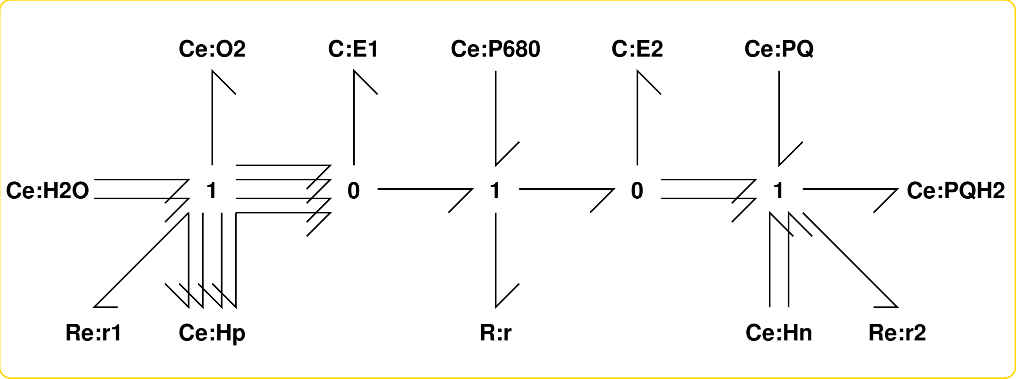

[30]:
```
imp.reload(sbg)
sbg.model('PII_abg.svg',convertCe=True,convertR=True,quiet=quiet)
import PII_abg
# imp.reload(PII_abg)
```

##### 6.1.1 Stoichiometry

[31]:
```
## Stoichiometry
sPII = st.stoich(PII_abg.model(),quiet=quiet)
chemostats = ['H2O','O2','PQ','PQH2','Hn','Hp','P680']
scPII = st.statify(sPII,chemostats=chemostats)
spPII = st.path(sPII,scPII)
## All reactions
disp.Latex(st.sprintrl(sPII,chemformula=chemformula))
```

[31]:

$$E_1 + P_{680} \xrightleftharpoons{r} E_2 \quad (21)$$

$$2\,H_2O \xrightleftharpoons{r_1} 4\,E_1 + 4\,Hp + O_2 \quad (22)$$

$$2\,E_2 + 2\,Hn + PQ \xrightleftharpoons{r_2} PQH_2 \quad (23)$$

##### 6.1.2 Pathway analysis with redox and pathway potentials

[32]:
```python
## Pathway analysis
disp.Latex(st.sprintrl(spPII,chemformula=chemformula))
```

[32]:

$$2\,H_2O + 4\,Hn + 4\,P_{680} + 2\,PQ \xrightleftharpoons{pr_1} 4\,Hp + O_2 + 2\,PQH_2 \quad (24)$$

[33]:
```python
## Compute net redox potential
RP_PII = (
            - redoxData.EpH('O2/2H2O',pH=pH_p)
            + V680
            + redoxData.EpH('PQ/PQH2',pH=pH_n)
        )

print(redoxData.EpH('O2/2H2O',pH=pH_p))
print(redoxData.E('P680+/P680*'))
print(redoxData.E7('PQ/PQH2'))
print(redoxData.EpH('PQ/PQH2',pH=pH_n))
#print(RP_PII)
print('RP_PII =',int(1000*RP_PII), 'mV')
```

```
1.0006212205771576
0.8
0
0.0
RP_PII = 822 mV
```

[34]:
```python
## Compute the reaction potential Phi
phi = phiData.phi_species(phi_redox,spPII['species'])
Phi_PII_ = -spPII['N'].T@phi
Phi_PII = Phi_PII_[0][0]
print(f'Phi_PII = {int(1000*Phi_PII)} mV')
print(f'Ratio = {(Phi_PII/RP_PII):.2f}')
```

```
Phi_PII = 3290 mV
Ratio = 4.00
```

[35]:
```python
## Photon contribution
n_phot=4
print(f"Photon contribution = {mV(n_phot*redoxData.E('P680+/P680*'))} mV")
```

```
Photon contribution = 3200 mV
```

[36]:
```python
## Potential per proton
n_prot = 4
print(f'Potential per proton = {mV(Phi_PII/n_prot)}')
```

```
Potential per proton = 823
```

#### 6.2 Complex Cyt – Cytochrome bf

[37]:
```
disp.SVG('Cyt_abg.svg')
```

[37]:

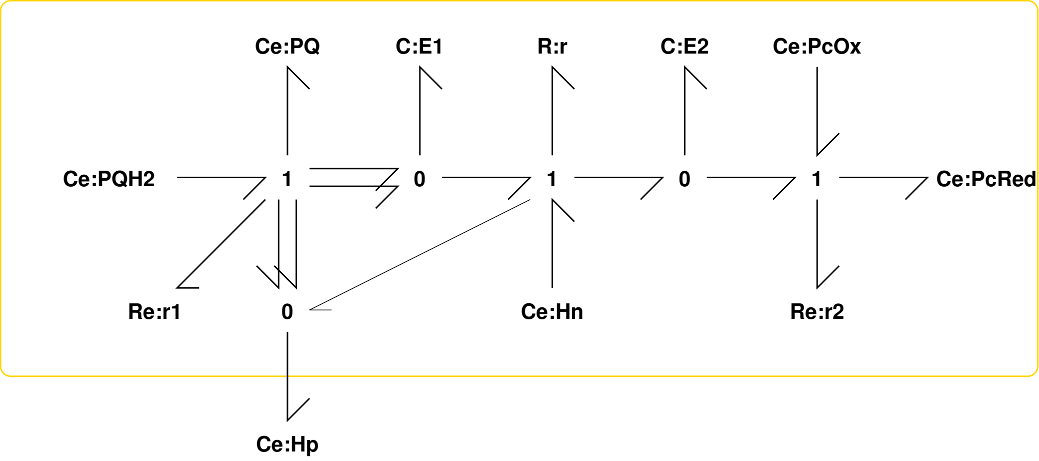

[38]:
```
sbg.model('Cyt_abg.svg',convertR=True,convertCe=True,quiet=quiet)
import Cyt_abg
# imp.reload(Cyt_abg)
```

##### 6.2.1 Stoichiometry

[39]:
```
## Stoichiometry
sCyt = st.stoich(Cyt_abg.model(),quiet=True)
chemostats = ['PQ','PQH2','PcOx','PcRed','Hp','Hn']
scCyt = st.statify(sCyt,chemostats=chemostats)
spCyt = st.path(sCyt,scCyt)
disp.Latex(st.sprintrl(sCyt,chemformula=chemformula))
```

[39]:

$$\mathtt{E_1} + \mathtt{Hn} \xrightleftharpoons[]{\mathtt{r}} \mathtt{E_2} + \mathtt{Hp} \tag{25}$$

$$\mathtt{PQH_2} \xrightleftharpoons[]{\mathtt{r_1}} 2\,\mathtt{E_1} + 2\,\mathtt{Hp} + \mathtt{PQ} \tag{26}$$

$$\mathtt{E_2} + \mathtt{PcOx} \xrightleftharpoons[]{\mathtt{r_2}} \mathtt{PcRed} \tag{27}$$

##### 6.2.2 Pathway analysis with redox and pathway potentials

[40]:
```
disp.Latex(st.sprintrl(spCyt,chemformula=chemformula))
```

[40]:

$$2\,\mathtt{Hn} + \mathtt{PQH_2} + 2\,\mathtt{PcOx} \xrightleftharpoons[]{\mathtt{pr_1}} 4\,\mathtt{Hp} + \mathtt{PQ} + 2\,\mathtt{PcRed} \tag{28}$$

[41]:
```
## Compute net redox potential
RP_Cyt = (- redoxData.EpH('PQ/PQH2',pH=pH_p)
          - redoxData.VpH(pH_p - pH_n)
```

```
         + redoxData.E('PcOx/PcRed')
        )

#print(RP_Cyt)
print('RP_Cyt =',redoxData.mV(RP_Cyt), 'mV')
```

```
RP_Cyt = 11 mV
```

[42]:
```
## Compute the reaction potential Phi
phi = phiData.phi_species(phi_redox,spCyt['species'])
Phi_Cyt_ = -spCyt['N'].T@phi
Phi_Cyt = Phi_Cyt_[0][0]
print('Phi_Cyt =',redoxData.mV(Phi_Cyt), 'mV')
print(f'Ratio = {(Phi_Cyt/RP_Cyt):0.2f}')
```

```
Phi_Cyt = 22 mV
Ratio = 2.00
```

##### 6.3 Complex PI – Photosystem I

[43]:
```
disp.SVG('PI_abg.svg')
```

[43]:

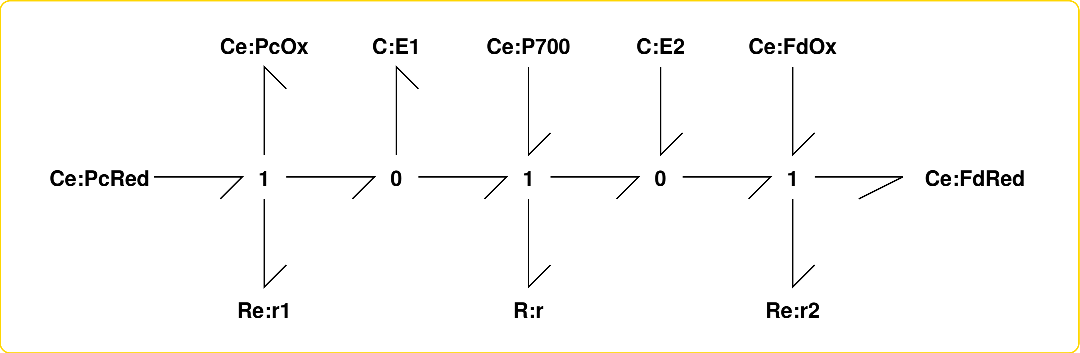

[44]:
```
sbg.model('PI_abg.svg',convertR=True,convertCe=True,quiet=True)
import PI_abg
# imp.reload(PI_abg)
```

###### 6.3.1 Stoichiometry

[45]:
```
## Stoichiometry
sPI = st.stoich(PI_abg.model(),quiet=quiet)
chemostats = ['PcOx','PcRed','FdOx','FdRed','P700']
scPI = st.statify(sPI,chemostats=chemostats)
spPI = st.path(sPI,scPI)
disp.Latex(st.sprintrl(sPI,chemformula=chemformula))
```

[45]:

$$E_1 + P_{700} \xrightleftharpoons{r} E_2 \quad (29)$$

$$PcRed \xrightleftharpoons{r_1} E_1 + PcOx \quad (30)$$

$$E_2 + FdOx \xrightleftharpoons{r_2} FdRed \quad (31)$$

##### 6.3.2 Pathway analysis with redox and pathway potentials

[46]:
```
disp.Latex(st.sprintrl(spPI,chemformula=chemformula))
```

[46]:

$$FdOx + P_{700} + PcRed \xrightleftharpoons{pr_1} FdRed + PcOx \quad (32)$$

[47]:
```
## Compute net redox potential
RP_PI = (
        - redoxData.E('PcOx/PcRed')
        + V700
        + redoxData.E('FdOx/FdRed')
        )

#print(RP_PI)
print('RP_PI =',redoxData.mV(RP_PI), 'mV')
```

```
RP_PI = 961 mV
```

[48]:
```
## Compute the reaction potential Phi
phi = phiData.phi_species(phi_redox,spPI['species'])
Phi_PI_ = -spPI['N'].T@phi
Phi_PI = Phi_PI_[0][0]
print('Phi_PI =',redoxData.mV(Phi_PI), 'mV')
print(f'Ratio = {(Phi_PI/RP_PI):0.2f}')
```

```
Phi_PI = 961 mV
Ratio = 1.00
```

#### 6.4 Complex Fer – Feredoxin-NADP reductase

[49]:
```
disp.SVG('Fer_abg.svg')
```

[49]:

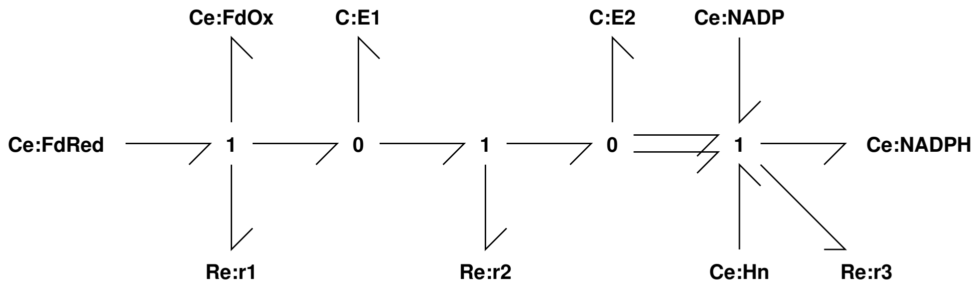

[50]:
```
sbg.model('Fer_abg.svg',convertR=True,convertCe=True,quiet=quiet)
import Fer_abg
# imp.reload(Fer_abg)
```

##### 6.4.1 Stoichiometry

[51]:
```
## Stoichiometry
sFer = st.stoich(Fer_abg.model(),quiet=True)
chemostats = ['FdRed','FdOx','NADP','NADPH','Hn']
scFer = st.stoich(Fer_abg.model(),chemostats=chemostats,quiet=True)
spFer = st.path(sFer,scFer)
disp.Latex(st.sprintrl(sFer,chemformula=chemformula))
```

[51]:

$$\mathtt{FdRed} \xrightleftharpoons{\mathtt{r}_1} \mathtt{E}_1 + \mathtt{FdOx} \tag{33}$$

$$\mathtt{E}_1 \xrightleftharpoons{\mathtt{r}_2} \mathtt{E}_2 \tag{34}$$

$$2\,\mathtt{E}_2 + \mathtt{Hn} + \mathtt{NADP} \xrightleftharpoons{\mathtt{r}_3} \mathtt{NADPH} \tag{35}$$

##### 6.4.2 Pathway analysis with redox and pathway potentials

[52]:
```
disp.Latex(st.sprintrl(spFer,chemformula=chemformula))
```

[52]:

$$2\,\mathtt{FdRed} + \mathtt{Hn} + \mathtt{NADP} \xrightleftharpoons{\mathtt{pr}_1} 2\,\mathtt{FdOx} + \mathtt{NADPH} \tag{36}$$

[53]:
```
## Compute net redox potential
RP_Fer = (
        - redoxData.E('FdOx/FdRed')
        + redoxData.EpH('NADP/NADPH',pH_n)
        )

#print(RP_Fer)
print('RP_Fer =',int(1000*RP_Fer), 'mV')
```

```
RP_Fer = 105 mV
```

[54]:
```
## Compute the reaction potential Phi
phi = phiData.phi_species(phi_redox,spFer['species'])
Phi_Fer_ = -spFer['N'].T@phi
Phi_Fer = Phi_Fer_[0][0]
print('Phi_Fer =',mV(Phi_Fer), 'mV')
print(f'Ratio = {(Phi_Fer/RP_Fer):0.2f}')
```

```
Phi_Fer = 212 mV
Ratio = 2.00
```

### 7 The Electron Transport Chain

- See section 7.1 of paper.

#### 7.1 Graphical description

[55]:
```
disp.SVG('ETCg_abg.svg')
```

[55]:

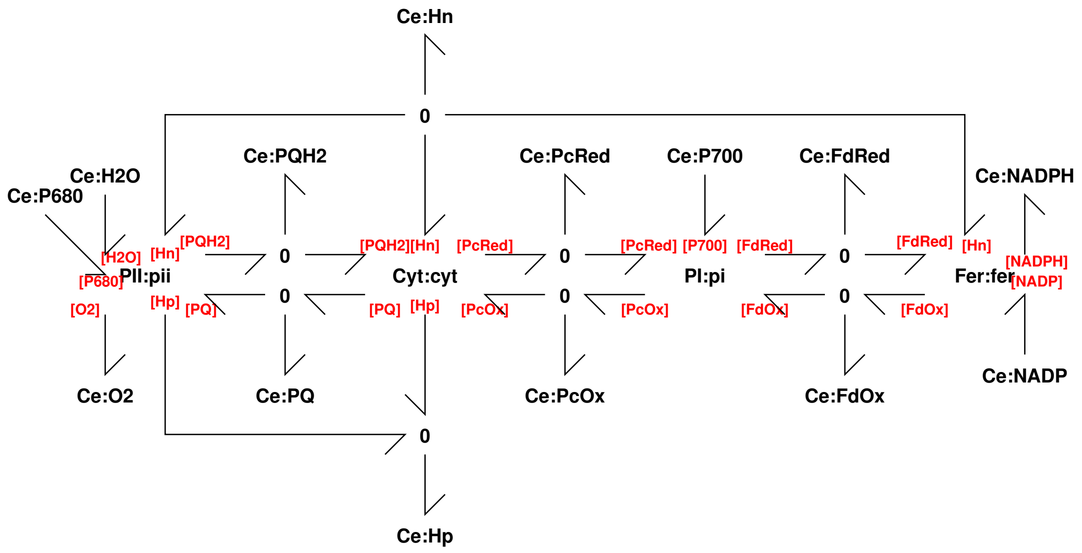

[56]:
```
sbg.model('ETCg_abg.svg',convertR=True,convertCe=True,quiet=quiet)
import ETCg_abg
imp.reload(ETCg_abg)
```

```
Creating subsystem: Cyt:cyt
Creating subsystem: Fer:fer
Creating subsystem: PI:pi
Creating subsystem: PII:pii
```

[56]:
```
<module 'ETCg_abg' from '/home/peterg/WORK/Dissemination/Edmund/SpecialIssue/
↪Exa
mples/Chloroplast/ETCg_abg.py'>
```

##### 7.1.1 Stoichiometry

[57]:
```
## Stoichiometry
sETCg = st.stoich(ETCg_abg.model(),quiet=True)
disp.Latex(st.sprintrl(sETCg,chemformula=chemformula))
```

[57]:

$$\mathtt{Hn} + \mathtt{cyt_{E1}} \xrightleftharpoons[]{\mathtt{cyt_r}} \mathtt{Hp} + \mathtt{cyt_{E2}} \tag{37}$$
$$\mathtt{PQH_2} \xrightleftharpoons[]{\mathtt{cyt_{r1}}} 2\,\mathtt{Hp} + \mathtt{PQ} + 2\,\mathtt{cyt_{E1}} \tag{38}$$
$$\mathtt{PcOx} + \mathtt{cyt_{E2}} \xrightleftharpoons[]{\mathtt{cyt_{r2}}} \mathtt{PcRed} \tag{39}$$
$$\mathtt{FdRed} \xrightleftharpoons[]{\mathtt{fer_{r1}}} \mathtt{FdOx} + \mathtt{fer_{E1}} \tag{40}$$
$$\mathtt{fer_{E1}} \xrightleftharpoons[]{\mathtt{fer_{r2}}} \mathtt{fer_{E2}} \tag{41}$$
$$\mathtt{Hn} + \mathtt{NADP} + 2\,\mathtt{fer_{E2}} \xrightleftharpoons[]{\mathtt{fer_{r3}}} \mathtt{NADPH} \tag{42}$$
$$\mathtt{P_{700}} + \mathtt{pi_{E1}} \xrightleftharpoons[]{\mathtt{pi_r}} \mathtt{pi_{E2}} \tag{43}$$
$$\mathtt{PcRed} \xrightleftharpoons[]{\mathtt{pi_{r1}}} \mathtt{PcOx} + \mathtt{pi_{E1}} \tag{44}$$
$$\mathtt{FdOx} + \mathtt{pi_{E2}} \xrightleftharpoons[]{\mathtt{pi_{r2}}} \mathtt{FdRed} \tag{45}$$
$$\mathtt{P_{680}} + \mathtt{pii_{E1}} \xrightleftharpoons[]{\mathtt{pii_r}} \mathtt{pii_{E2}} \tag{46}$$

$$2\,\mathtt{H_2O} \xrightleftharpoons[]{\mathtt{pii_{r1}}} 4\,\mathtt{Hp} + \mathtt{O_2} + 4\,\mathtt{pii_{E1}} \tag{47}$$
$$2\,\mathtt{Hn} + \mathtt{PQ} + 2\,\mathtt{pii_{E2}} \xrightleftharpoons[]{\mathtt{pii_{r2}}} \mathtt{PQH_2} \tag{48}$$

##### 7.1.2 Pathway analysis with redox and pathway potentials

[58]:
```
chemostats = ['P680','P700','H2O','O2','Hn','Hp','NADP','NADPH']
scETCg = st.stoich(ETCg_abg.model(),chemostats=chemostats,quiet=True)
spETCg = st.path(sETCg,scETCg)
disp.Latex(st.sprintrl(spETCg,chemformula=chemformula,all=True))
```

[58]:

$$2\,\mathtt{H_2O} + 10\,\mathtt{Hn} + 2\,\mathtt{NADP} + 4\,\mathtt{P_{680}} + 4\,\mathtt{P_{700}} \xrightleftharpoons[]{\mathtt{pr_1}} 12\,\mathtt{Hp} + 2\,\mathtt{NADPH} + \mathtt{O_2} \tag{49}$$

[59]:
```
## Compute the reaction potential Phi
species = spETCg['species']
print(species)
phi = phiData.phi_species(phi_redox,species)
Phi_ = -spETCg['N'].T@phi
Phi= Phi_[0][0]
print('Phi =',mV(Phi), 'mV')
```

```
['FdRed', 'H2O', 'Hn', 'Hp', 'NADP', 'NADPH', 'O2', 'P680', 'P700', 'PQH2',
'PcRed']
Phi = 7603 mV
```

[60]:
```
## Flatten
stbg.model(sETCg,filename='ETCg_abg')
import ETCg_abg
imp.reload(ETCg_abg)
```

[60]:

```
<module 'ETCg_abg' from '/home/peterg/WORK/Dissemination/Edmund/SpecialIssue/
↪Exa
mples/Chloroplast/ETCg_abg.py'>
```

##### 7.2 Computational description

The overall model is described a bond graph tools file:

[61]:

```
## File ETC_abg.py

## Import the modules
import PII_abg
import Cyt_abg
import PI_abg
import Fer_abg

## Create the model using BondGraphTools
def model():
    """
    Model of chloroplast electron transport chain
    """

    ETC = bgt.new(name='ETC')   # Create system
    PII = PII_abg.model()
    Cyt = Cyt_abg.model()
    PI = PI_abg.model()
    Fer = Fer_abg.model()
    bgt.add(ETC,PII,Cyt,PI,Fer)

    return ETC
```

###### 7.2.1 Unify species in model using mbg.unify()

[62]:

```
# import ETC_abg
ETC = model()
common = ['PQ','PQH2','PcOx','PcRed','FdOx', 'FdRed','Hn','Hp']
mbg.unify(ETC,common,quiet=quiet)
ss = st.stoich(ETC,quiet=quiet)
disp.Latex(st.sprintrl(ss,chemformula=chemformula))
```

[62]:

$$E_1 + P_{680} \xrightleftharpoons{r} E_2 \tag{50}$$
$$2\,H_2O \xrightleftharpoons{r_1} 4\,E_1 + O_2 + 4\,Hp \tag{51}$$
$$2\,E_2 + PQ + 2\,Hn \xrightleftharpoons{r_2} PQH_2 \tag{52}$$
$$E_1 + Hn \xrightleftharpoons{r} E_2 + Hp \tag{53}$$
$$PQH_2 \xrightleftharpoons{r_1} 2\,E_1 + PQ + 2\,Hp \tag{54}$$
$$E_2 + PcOx \xrightleftharpoons{r_2} PcRed \tag{55}$$
$$E_1 + P_{700} \xrightleftharpoons{r} E_2 \tag{56}$$
$$PcRed \xrightleftharpoons{r_1} E_1 + PcOx \tag{57}$$
$$E_2 + FdOx \xrightleftharpoons{r_2} FdRed \tag{58}$$
$$FdRed \xrightleftharpoons{r_1} E_1 + FdOx \tag{59}$$

$$E_1 \xrightleftharpoons{r_2} E_2 \tag{60}$$
$$2\,E_2 + NADP + Hn \xrightleftharpoons{r_3} NADPH \tag{61}$$

### 8 Chloroplast

- See section 7.1 of paper.

#### 8.1 Modular model

[63]:
```
disp.SVG('Chloroplast_abg.svg')
```

[63]:

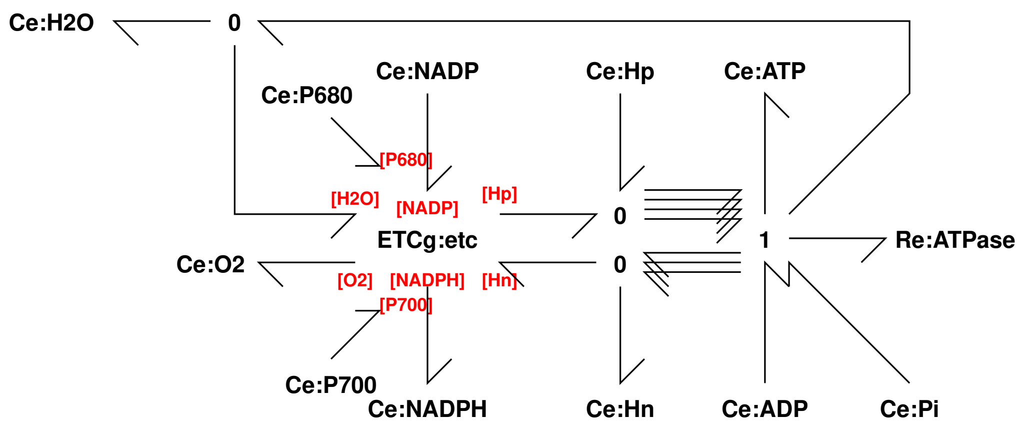

[64]:
```
sbg.model('Chloroplast_abg.svg',convertR=True,convertCe=True,quiet=quiet)
import Chloroplast_abg
imp.reload(Chloroplast_abg)
```

```
Creating subsystem: ETCg:etc
```

[64]: <module 'Chloroplast_abg' from '/home/peterg/WORK/Dissemination/Edmund/
↪SpecialIs
sue/Examples/Chloroplast/Chloroplast_abg.py'>

#### 8.2 Generate equations

[65]:
```
## Stoichiometry
sChloroplast = st.stoich(Chloroplast_abg.model(),quiet=True)
disp.Latex(st.sprintrl(sChloroplast,chemformula=chemformula))
```

[65]:

$$Hn + etc_cyt_{E1} \xrightleftharpoons{etc_cyt_r} Hp + etc_cyt_{E2} \tag{62}$$

$$etc_pQH_2 \xrightleftharpoons{etc_cyt_{r1}} 2\,Hp + etc_pQ + 2\,etc_cyt_{E1} \tag{63}$$

$$etc_pcOx + etc_cyt_{E2} \xrightleftharpoons{etc_cyt_{r2}} etc_pcRed \tag{64}$$

$$etc_FdRed \xrightleftharpoons{etc_fer_{r1}} etc_FdOx + etc_fer_{E1} \tag{65}$$

$$etc_fer_{E1} \xrightleftharpoons{etc_fer_{r2}} etc_fer_{E2} \tag{66}$$

$$Hn + NADP + 2\,etc_fer_{E2} \xrightleftharpoons{etc_fer_{r3}} NADPH \tag{67}$$

$$P_{700} + etc_pi_{E1} \xrightleftharpoons{etc_pi_r} etc_pi_{E2} \tag{68}$$

$$etc_pcRed \xrightleftharpoons{etc_pi_{r1}} etc_pcOx + etc_pi_{E1} \tag{69}$$

$$etc_FdOx + etc_pi_{E2} \xrightleftharpoons{etc_pi_{r2}} etc_FdRed \tag{70}$$

$$P_{680} + etc_pii_{E1} \xrightleftharpoons{etc_pii_r} etc_pii_{E2} \tag{71}$$

$$2\,H_2O \xrightleftharpoons{etc_pii_{r1}} 4\,Hp + O_2 + 4\,etc_pii_{E1} \tag{72}$$

$$2\,Hn + etc_pQ + 2\,etc_pii_{E2} \xrightleftharpoons{etc_pii_{r2}} etc_pQH_2 \tag{73}$$

$$ADP + 4\,Hp + Pi \xrightleftharpoons{ATPase} ATP + H_2O + 3\,Hn \tag{74}$$

#### 8.3 Pathway analysis

[66]:
```
# imp.reload(st)
chemostats = ['P680','P700','H2O','O2','Hn','NADP','NADPH','ATP','ADP','Pi']
scChloroplast = st.stoich(Chloroplast_abg.
↪model(),chemostats=chemostats,quiet=True)
spChloroplast = st.path(sChloroplast,scChloroplast)
```

##### 8.3.1 Pathway reaction

[67]:
```
disp.Latex(st.sprintrl(spChloroplast,chemformula=chemformula,all=True))
```

[67]:

$$3\,ADP + Hn + 2\,NADP + 4\,P_{680} + 4\,P_{700} + 3\,Pi \xrightleftharpoons{pr_1} 3\,ATP + H_2O + 2\,NADPH + O_2 \tag{75}$$

#### 8.4 Efficiency from estimated $\phi$

[68]:
```
## Compute the reaction potential Phi
species = spChloroplast['species']
# species = chemostats
# print(species)

phi = phiData.phi_species(phi_redox,species)

Phi_ = -spChloroplast['N'].T@phi
Phi= Phi_[0][0]
print(f'Phi = {Phi:0.2f} V')

## Photo input
# print(phi_redox['P680'] , phi_redox['P700'])
Phi_phot = 4*(phi_redox['P680'] + phi_redox['P700'])
print(f'Phi_phot = {Phi_phot:0.2f} V')

##Outputs
Phi_out = Phi_phot - Phi
print(f'Phi_out = {Phi_out:0.2f} V')

## Efficiency
efficiency = Phi_out/Phi_phot
print(f'Efficiency = {(efficiency*100):.1f} %')
```

```
Phi = 8.73 V
Phi_phot = 14.38 V
Phi_out = 5.65 V
Efficiency = 39.3 %
```

[69]:
```
## Compute the reaction potential Phi but with no photons
phi = phiData.phi_species(phi_redox_0,species)

Phi_ = -spChloroplast['N'].T@phi
Phi_out = -Phi_[0][0]
print(f'Phi_out = {Phi_out:0.2f} V')

## Photo input
# print(phi_redox['P680'] , phi_redox['P700'])
Phi_phot = 4*(phi_redox['P680'] + phi_redox['P700'])
print(f'Phi_phot = {Phi_phot:0.2f} V')

## Efficiency
efficiency = Phi_out/Phi_phot
print(f'Efficiency = {(efficiency*100):.1f} %')
```

```
Phi_out = 5.65 V
Phi_phot = 14.38 V
Efficiency = 39.3 %
```

#### 8.5 Efficiency by direct calculation

- See section 7.2 of paper.

[70]:
```
def getPhi(name):
    redox_data = redoxData.data()
    n = redox_data[name]['electrons']
    E = redox_data[name]['E7']
    Phi = n*E
    print(f'{name}:\t E = {E} V;\t n = {n};\t Phi = {Phi} V')
    return Phi
```

[71]:
```
## Photons
Phi_phot = 4*(phi_redox['P680'] + phi_redox['P700'])

print(f'Phi_phot = {Phi_phot:0.2f} V')
## NADPH
Phi_1 = getPhi('NADP/NADPH')

## O2
Phi_2 = getPhi('O2/2H2O')

## ATP Hydrolysis
print(f'Hydrolysis:\t Phi = {Phi_Hyd:0.3f}')
```

```
Phi_phot = 14.38 V
NADP/NADPH:      E = -0.324 V;   n = 2;  Phi = -0.648 V
O2/2H2O:         E = 0.816 V;    n = 4;  Phi = 3.264 V
Hydrolysis:      Phi = 0.363
```

[72]:
```
## All
Phi_chem = -2*Phi_1 + Phi_2 + 3*Phi_Hyd
print(f'Phi_chem = {Phi_chem:0.2f}')
```

```
Phi_chem = 5.65
```

[73]:
```
## Efficiency
efficiency = Phi_chem/Phi_phot
print(f'Efficiency = {(efficiency*100):.1f} %')
```

```
Efficiency = 39.3 %
```

[ ]:
```

```
